# High-Throughput Single-Molecule R-loop Footprinting Reveals Principles of R-loop Formation

**DOI:** 10.1101/640094

**Authors:** Maika Malig, Stella R. Hartono, Jenna M. Giafaglione, Lionel A. Sanz, Frederic Chedin

**Affiliations:** Department of Molecular and Cellular Biology and Genome Center, University of California, Davis, Davis, CA 95616; Integrative Genetics and Genomics graduate group, University of California Davis, Davis, CA 95616; Molecular Biology Interdepartmental Doctoral Program, University of California Los Angeles, Los Angeles, CA 90095-1570

**Keywords:** R-loops, DNA:RNA immunoprecipitation, SMRT sequencing, non-denaturing bisulfite conversion, DNA topology, S9.6 antibody

## Abstract

R-loops are a prevalent class of non-B DNA structures that form during transcription upon reannealing of the nascent RNA to the template DNA strand. R-loops have been profiled using the S9.6 antibody to immunoprecipitate DNA:RNA hybrids. S9.6-based DNA:RNA immunoprecipitation (DRIP) techniques revealed that R-loops form dynamically over conserved genic hotspots. We developed an orthogonal profiling methodology that queries R-loops via the presence of long stretches of single-stranded DNA on the looped-out strand. Non-denaturing sodium bisulfite treatment catalyzes the conversion of unpaired cytosines to uracils, creating permanent genetic tags for the position of an R-loop. Long read, single-molecule PacBio sequencing allows the identification of R-loop ‘footprints’ at near nucleotide resolution in a strand-specific manner on single DNA molecules and at ultra-deep coverage. Single-molecule R-loop footprinting (SMRF-seq) revealed a strong agreement between S9.6-and bisulfite-based R-loop mapping and confirmed that R-loops form from unspliced transcripts over genic hotspots. Using the largest single-molecule R-loop dataset to date, we show that individual R-loops generate overlapping sets of molecular clusters that pile-up through larger R-loop-prone zones. SMRF-seq further established that R-loop distribution patterns are driven by both intrinsic DNA sequence features and DNA topological constraints, revealing the principles of R-loop formation.

## INTRODUCTION

R-loops are three-stranded nucleic acid structures that primarily form during transcription as a result of the nascent RNA annealing to the template DNA strand [1]. The formation of this RNA:DNA hybrid causes the non-template DNA strand to loop out in a single-stranded state. R-loops have been comprehensively mapped at genome scale [2, 3] using the S9.6 antibody to immunoprecipitate the RNA:DNA hybrid moiety of the structure [4, 5]. DNA:RNA Immunoprecipitation (DRIP) based studies revealed that R-loops are prevalent and dynamic non-B DNA structures that form over specific and conserved genic hotspots during transcription [3]. An accumulating body of data has linked R-loops to both physiological and pathological outcomes [6, 7]. On one hand, R-loops have been linked with efficient transcription termination [3, 8], chromatin patterning [3, 9], and the regulation of gene expression [10, 11]. On the other hand, deregulated R-loop formation has been implicated in genome instability arising from transcription-replication conflicts caused by dysfunctions in a variety of nuclear factors, including RNA processing enzymes [1, 12-16]. Deregulated R-loop formation has also been implicated in a number of human diseases, including certain cancers and neurological disorders [17].

While DRIP-based methods provide robust and specific information on the global distribution of R-loops in a genome, they suffer from a few important limitations. First, DRIP-based methods only provide a population average representation of R-loop distribution; the exact positions or lengths of individual R-loops cannot be deduced from the data. Developing single-molecule R-loop profiling methods will allow us to elucidate the precise mechanisms underlying R-loop formation and stability and may provide insights into the structural features that distinguish normal from pathological R-loops. Second, DRIP-based methods are exclusively dependent on the S9.6 antibody. While S9.6 possesses sub-nanomolar affinity to RNA:DNA hybrids, several studies have highlighted that it possesses a significant residual affinity for double-stranded RNA (dsRNA) [4, 18]. This affinity complicates the use of S9.6 in R-loop profiling when dsRNA are produced [18] even though adequate ribonuclease pretreatments can be used to mitigate these issues. Due to the rising interest in R-loop biology, the development of orthogonal R-loop profiling methodologies that are S9.6 independent is desirable. The development of a method that enables the identification of R-loop positions at near nucleotide resolution in a strand-specific manner on single DNA molecules and at high coverage independently of S9.6 enrichment would set a strong benchmark for the field.

Prior to the development of S9.6-based DRIP methods [2, 8], endogenous R-loops were queried not through their RNA:DNA hybrid moiety but through their association with long stretches of single-stranded DNA (ssDNA) on the displaced non-template strand [19]. This method took advantage of the fact that sodium bisulfite is an exquisitely sensitive probe of ssDNA and efficiently triggers the deamination of exposed cytosines (C) to uracils (U) [20]. When nucleic acids extracted from cells are treated with sodium bisulfite in a non-denaturing manner, intrinsically single-stranded regions undergo C to U conversion. These patches of uracils serve as permanent genetic tags for the position of a given R-loop (Figure S1). After PCR amplification, uracils are converted to thymines (T) and the resulting amplicons can be cloned and sequenced individually using Sanger sequencing. Mapping patches of C to T conversion permits single-molecule R-loop footprinting [2, 19, 21]. As with S9.6-based methods, pre-treatment of nucleic acids with purified Ribonuclease H (RNase H), an enzyme that specifically degrades RNA strands in the context of RNA:DNA hybrids [22], serves as a robust specificity control. The application of this method to R-loops induced at class-switch regions in murine B cells has provided numerous insights into the mechanisms of R-loop formation in mammalian genomes [19, 21, 23, 24]. One drawback of this method, however, is that it suffers from low throughput and requires laborious, time-consuming and cost-intensive cloning, sequencing, and data analysis steps. Here we present a novel adaption of R-loop footprinting that permits single-molecule R-loop detection at near nucleotide resolution in a strand-specific manner on long amplicons and at ultra-high coverage. This method and its accompanying computational analysis and data visualization pipeline enables deep, cost-effective, R-loop profiling at a range of genomic loci, under any condition and in any genome.

## MATERIAL AND METHODS

### Non-denaturing Single-Molecule R-loop Footprinting coupled to PacBio sequencing

#### Cell culture

NTERA-2 cells (ATCC® CRL-1973) were used for all footprinting experiments and grown under standard conditions in a humidified incubator at 5% CO_2_ in DMEM high glucose media supplemented with 10% FBS. Cells were grown to 80-90% confluence and split to 50% in new media and harvested 16 hours later.

#### Nucleic acid isolation, fragmentation and immunoprecipitation

DNA extraction was done as described for standard DRIP-seq protocol [2] except that the proteinase K incubation was shortened to 2 hours instead of overnight. Restriction enzymes (REs) were selected to fragment genomic DNA around target regions so that fragments of no more than 15 kb were generated. We used 80-120 units total of REs and incubated overnight at 37°C followed by a DNA purification step using 1X AMPure beads. This digested template was either sodium bisulfite treated directly to footprint R-loops or after immunoprecipitation using the anti-RNA:DNA hybrid S9.6 antibody [5] according to the standard DRIP protocol.

#### Non-denaturing bisulfite conversion

Samples were treated with sodium bisulfite according to the Zymo EZ DNA Methylation-Lightning kit (PN D5030) following the manufacturer’s instructions except for a few critical modifications to permit the probing of intrinsically single-stranded cytosines under non-denaturing conditions. In brief, the DNA was not denatured prior to bisulfite treatment and treatment with the conversion reagent was performed at 37°C for 2 hours with gentle rotation. Desulfonation and sample recovery were done as instructed. As a positive control, a linearized denatured plasmid was spiked-in with genomic samples and treated accordingly. All recovered DNA molecules were heavily converted and the overall C to T conversion efficiency was 86% (Figure S10). This indicates that bisulfite treatment under these conditions was efficient and suitable for R-loop detection. Direct non-denaturing bisulfite conversion was performed according to procedure B in the Zymo EZ DNA Methylation-Direct kit (PN D5020) with a few modifications. Briefly, samples containing approximately 30,000 cells were pelleted and resuspended in 1 mL 1X DPBS, lysed with 1X final concentration of M-Digestion buffer, and incubated with Proteinase K (included in the kit) for 25 minutes at 37°C. The digested sample was then directly treated with sodium bisulfite at 37°C for 3.5 hours with gentle rotation. The following steps were performed as per manufacturer’s instructions.

#### Site-specific PCR amplification

We comprehensively surveyed R-loop forming gene regions targeting the promoter, terminal, or gene body of selected genes using available DRIPc-seq data. Amplicons ranging from 2-4 kb were selected to allow sufficient coverage for each locus and efficient PacBio sequencing. Primers were designed using Primer3 (bioinfo.ut.ee/primer3-0.4.0) (Table S2). PCR reactions using ThermoFisher PhusionU DNA polymerase (ThermoFisher PN F555S) were optimized to produce long-range, high fidelity single-band products (conditions available upon request). Standard PCR reaction included 1X PCR buffer, 0.2 mM dNTPs, 0.8 μM each forward and reverse primers, 5-30 ng DNA template, 0.02 U/ μL Phusion U DNA polymerase, 1M Betaine (optional), and PCR grade water to a 30 μL final volume. PCR cycling was as follows: 1) Initial denaturing at 98°C for 30 seconds; 2) 30-35 cycles of: 2a. denature at 98°C for 10 seconds, 2b. anneal at optimized temperature for 30 seconds, 2c. extend at 72°C for 2.5 minutes; 3) Final extension at 72°C for 5 minutes; 4) Hold at 4°C infinitely. All PCR products were purified using 1X Ampure beads.

#### SMRTbell library construction

We used the PacBio RSII system to achieve long-read, single-molecule resolution sequencing of R-loop footprints. We generated libraries by pooling non-overlapping amplicons (less than 20 products per run) adding equal amounts for each. Starting with 1-2 μg of PCR products, pooled samples were concentrated using 1X Ampure bead wash. Libraries were built following the “Procedure & Checklist – 2 kb Template Preparation and Sequencing” protocol (PN 001-143-835-08) from PacBio with a few modifications. No prior DNA damage repair step was done. AMPure bead wash steps were done using 0.8X concentration. Ligation was done for 1 hour at 25°C. SMRTbell libraries were quantified and size confirmation done by either by gel electrophoresis or Agilent Genomic’s 2100 Bioanalyzer. Libraries were sequenced on a PacBio RSII instrument with 6-hour movie times.

### R-loop Profiling using DRIPc-seq

DRIPc-seq was applied to NTERA-2 cells as described [3] except a Ribonuclease A pre-treatment was applied to the extracted nucleic acids prior to S9.6 immunoprecipitation.

### Computational data processing

#### Circular consensus sequence (CCS) generation

Subreads of read quality at least 90% were further processed into CCS using PacBio SMRT Analysis pipeline (ConsensusTools.sh) with a minimum pass filter of 3.

#### Duplicate read removal

We used dedupe2.sh from package BBMap V37.90 (*https://sourceforge.net/projects/bbmap/*) with default parameters except for mid=98, nam=4, k=31, and e=30. An average of 43% reads (254,951 out of 592,444 total CCS reads) were removed as duplicates in the combined datasets.

#### Gargamel computational pipeline

The Gargamel pipeline (available at https://github.com/srhartono/footLoop) allows user to map reads, assign strand, call single molecule R-loop footprints (SMRFs) as peaks of C to T conversion, perform clustering on SMRFs, and visualize the data.

#### Read mapping

Reads were mapped to the hg19 human genome reference focusing on their respective amplicon regions using Bismark v0.13.1 [26] including a 10 bp buffer off their beginning and end positions. Bismark default settings were used except for a slightly relaxed minimum score threshold (--score_min of L,0,-0.3 instead of L,0,-0.2). Truncated reads shorter than 95% of their expected length were discarded. Altogether, the stringent requirements imposed for circular consensus, duplicate removal, mapping, and size, ensured the selection of very high-quality reads.

#### Strand assignment

For each read, we assigned strand based on their conversion patterns. Reads with insufficient conversions (C->T < 6 and G->A < 6) could not be assigned a strand. Likewise, if the number of C->T conversions was within +/-10% of G->A conversions on a read, then the strandedness was considered as unknown. Such reads represented less than ∼5% of the total pool. Otherwise, reads with predominant C to T or G to A conversions were assigned as non-template or template strand, respectively. Ambiguous regions carrying indels due to PacBio sequencing errors were masked (including a 5 bp buffer around the indel) so as not to distort the conversion frequency calculation. Some individual loci consistently resulted in more reads mapping to one strand than to the other, regardless of which strand carried R-loop footprints, most likely reflecting PCR strand biases. Considering all tested loci together, reads were approximately equally distributed to each strand.

#### Peak calling

A threshold-based sliding window method was used to call tracts of C to T conversion referred to as R-loop peaks. The windows spanned 20 consecutive cytosines and were moved across each read one cytosine at a time. For each window, we calculated the C to T conversion frequency and called a window R-loop positive if a minimum of 55% cytosines were converted. Positive windows were further extended if neighboring windows also satisfied the 55% threshold. Upon encountering a window with conversion frequency less than the threshold, the peak was terminated and its boundaries recorded. In this study, we imposed that each positive peak should be at least 100 bp in length. We tested a combination of window sizes (10, 15 and 20 cytosines), conversion thresholds (35%, 55%, 65%, 70%, 75%, and 80%), and minimal peak sizes (30 and 100 bp); the results were qualitatively similar except that more positive peaks were recovered for less stringent conditions. A window size of 20 cytosines, minimal C to T conversion of 55%, and a minimal length of 100 bp permitted a good combination of specificity and sensitivity. For samples directly treated with bisulfite after a short Proteinase K digestion, peak calling thresholds were modified to 35% conversion frequency with 30 bp minimum peak length.

#### Clustering

For each gene, R-loop peaks were clustered using their start and stop coordinates using k-means clustering. K was determined automatically by minimizing intra-cluster distances, iterating until minimum within-group distance for each cluster was at most three.

#### Reproducibility and location analysis

For each locus, we combined reads from all replicates and generated clusters as described above. For each biological replicate, we then quantified the distribution of that replicate’s reads across these predefined clusters. If independent replicates are sampling from the same overall biological distribution, the expectation is that read distributions across replicates will be similar. Reproducibility was therefore measured by calculating Pearson correlation coefficients applied to the read distribution across clusters between replicates. As a control, we shuffled the position of each read around the amplicon region (keeping length information intact) and assigned the shuffled reads to the same set of predefined clusters. A shuffled read was considered to belong to a specific cluster if its start and end positions fell within +/-100 bp of the mean start and end of all experimental reads in that cluster. Reads that could not be assigned to any experimentally determined cluster were assigned to an extra “shuffled” cluster. The Pearson correlation coefficients between experimental and shuffled reads were calculated across clusters as described above.

#### Identification of template-strand ssDNA patches at the edge of R-loop clusters

C to T conversion frequencies were measured on the template strand over a 20 bp window around the edges of R-loop footprint clusters. Molecules with conversion frequencies >5% were selected. To determine if these conversion events were a product of random DNA breathing or reflected the presence of a bisulfite-susceptible ssDNA patch on certain molecules, we subtracted background C to T conversion frequencies for all non-template strand molecules. This was performed either by subtracting the average background C to T conversion frequency across the entire amplicon or the average C to T conversion frequency for each position along the amplicon, with similar results. As a negative control, we determined if randomly assigned clusters outside of actual footprint regions would show C to T conversion patches at their edges. Shuffled clusters were not associated with significant template-strand ssDNA patches.

## RESULTS

### Non-denaturing bisulfite R-loop footprinting coupled with high-throughput single-molecule sequencing

Non-denaturing bisulfite treatment allows R-loop mapping by promoting the deamination of intrinsically single-stranded cytosines to uracils on the looped-out DNA strand of an R-loop [19]. To achieve orders of magnitude higher throughput than possible using conventional sequencing methods, Pacific Biosciences (PacBio) libraries were built directly from PCR amplicons and sequenced using single-molecule, real-time sequencing (SMRT-seq; Figure S1). SMRT-seq is well suited for this purpose as it allows sequencing through kilobases-long GC-rich amplicons [25] and delivers high-quality sequencing reads on 50,000 single DNA molecules per SMRT cell on a PacBio RSII instrument. Given this throughput, several genomic regions can be tested for R-loop formation in one SMRT cell without any prior enrichment. In the resulting method, <u>s</u>ingle-<u>m</u>olecule <u>R</u>-loop footprinting coupled with PacBio sequencing (SMRF-seq), C to T conversion patterns expected from true R-loops should satisfy the following predictions: 1) R-loops can be detected upon amplification of regions with native primers without any prior enrichment with the S9.6 antibody; 2) R-loop C to T conversion tracts can only be observed on the looped out strand, not on the RNA-paired strand; and 3) C to T conversion tracts on the looped-out strand are lost upon RNase H treatment of the genomic DNA prior to bisulfite treatment. Accurately extracting C to T conversion information from SMRF-seq runs required the development of a new computational pipeline. In brief, high-quality circular consensus sequences (CCS) were built and mapped against reference amplicons using the bisulfite mapping-enabled pipeline Bismark [26]. Strand assignments were based on the patterns of C to T or G to A conversions as described [19] (Figure S1). Long, R-loop-associated C to T conversion footprints were then called using a threshold-based method in which a sliding window encompassing 20 consecutive cytosines were analyzed for their conversion status and a minimum user-imposed conversion threshold was applied (see Methods for details). The imposition of a minimal window size facilitated our ability to distinguish R-loop footprints from small-scale events of DNA breathing and from PacBio sequencing errors. Note that patterns of bisulfite conversion were only analyzed for non-CpG dinucleotides since methylation of CpG sites blurs our ability to interrogate the ssDNA status of these sites. Non-CpG methylation is present only at very low levels in the human Ntera2 cell line used here [3].

### S9.6-independent, strand-specific R-loop footprints in the human genome

As a proof of concept, we investigated R-loop formation at five loci (*FUS, RPL13A, SNRNP70, RPS24* and *PIN4*) that exhibited prominent R-loop peaks previously identified using S9.6-based methods^3^. *PIN4* represents an example of a GC-skewed promoter, while *FUS, RPL13A,* and *SNRNP70* represent examples of gene body / early terminal R-loops and *RPS24* represents a terminal R-loop peak. We used SMRF-seq to analyze these regions in the absence of any S9.6 pre-enrichment.

In all cases, prominent R-loop footprints were observed specifically on the non-template strand for transcription. Using *FUS* as a representative example, a total of 226 footprints were called on this strand out of 5,111 molecules sequenced (Figure 1). Thus the proportion of R-loop-carrying molecules was 4.42% which is in close agreement with the range of R-loop frequencies measured by DRIP-qPCR [3]. Only three footprints were detected on the template strand out of 14,489 molecules sequenced, showing that bisulfite-reactive ssDNA was essentially confined to the non-template strand. Importantly, the collection of footprints observed at *FUS* was in strong agreement with bulk S9.6-based DRIPc-seq profiles obtained independently from the same cells. SMRF-seq delineated three regions where footprints were more likely to pile up; these three regions matched remarkably well with the corresponding sub-peaks in DRIPc-seq data (Figure 1). As a result, aggregate bisulfite conversion profiles over R-loop peaks were highly congruent with DRIPc-seq. Similar results were observed for all other tested loci (Figure S2). This demonstrates that strand-specific R-loop footprints can be readily recovered at a range of R-loop-prone loci in the human genome using SMRF-seq. Furthermore, it shows that non-denaturing bisulfite footprinting-based and orthogonal S9.6-based approaches report on R-loop formation with strong agreement in terms of location and strandedness. Unlike population-based approaches, however, SMRF-seq allows us to peer into the collection of individual R-loops that together give rise to R-loop hotspots.

**Figure 1:**
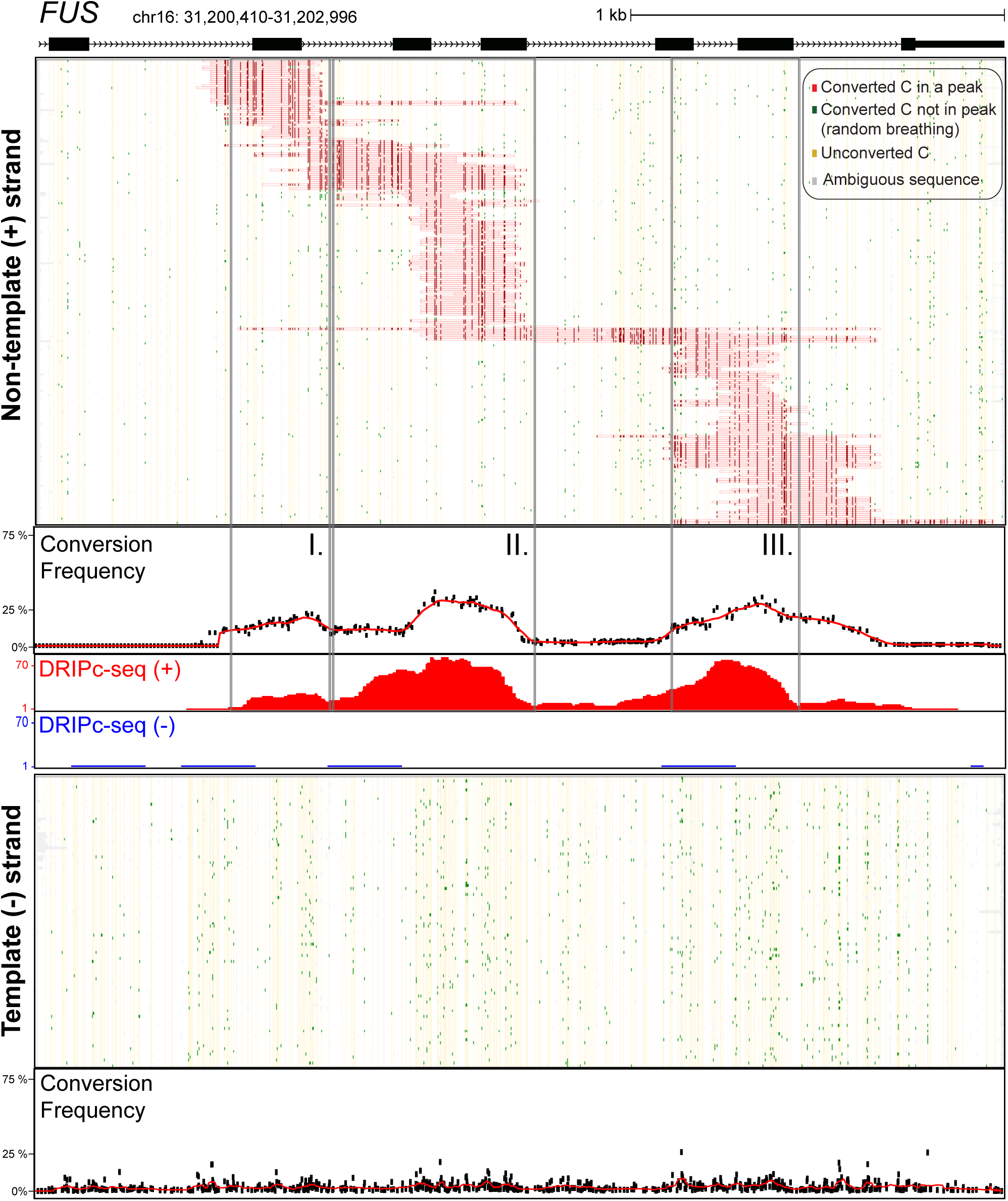
Single-molecule R-loop footprints at the human *FUS* locus. The structure of *FUS* over the amplicon is shown at top with exons as boxes. The coordinates (hg19) of the amplicon are shown along with a scale bar. 226 independent molecules carrying R-loop footprints on the non-template strand are shown at top. Each horizontal line corresponds to one DNA molecule. The status of each cytosine along the sequence is color-coded with footprints highlighted in red (see inset for color codes). These footprints were obtained using SMRF-seq with native PCR primers independently of any S9.6 enrichment. The aggregate C to T conversion frequency calculated over these footprints is shown below along with the bulk S9.6-based R-loop signals obtained independently from DRIPc-seq (red indicates signal on the (+) strand; blue on the (-) strand). Vertical boxes highlight three regions where R-loop footprints tend to pile up. 100 independent molecules recovered from the template strand were randomly sampled from the 14,489 sequenced and are displayed below along with the aggregate C to T conversion frequency for that strand.

### DRIP enriches but does not disturb genuine R-loop footprints

Given the overall agreement between aggregate SMRF-seq data and DRIPc-seq profiles, we asked whether the location or relative distribution of footprints would be modified by prior S9.6 enrichment. We therefore conducted DRIP followed by SMRF-seq and analyzed the same five loci previously described. As expected, the proportion of footprint-containing molecules was five to tenfold higher after DRIP enrichment at most loci (Table 1). Post-DRIP footprints were consistent with those observed without enrichment: they were strand-specific and their distribution in terms of aggregate profiles was consistent with and without S9.6 enrichment as shown for *RPS24* and *FUS* (Figure 2; Pearson’s R were 0.99 and 0.94, respectively) and other loci (Figure S3). When analyzed in terms of location and lengths, R-loop footprints with and without S9.6 enrichment were again mostly consistent across loci although minor differences were noted for individual loci (Figure S4). For the *SNRPN70* locus, one class of R-loops appeared differentially represented with and without S9.6 enrichment leading to a discrepancy on location and lengths, but it is unclear if this difference was biologically significant or resulted from possible under-sampling of the R-loop distribution. Overall, S9.6-based DRIP allowed the efficient enrichment of footprints without significantly disturbing their properties.

**Table 1:** Summary of SMRF-seq data comparing footprinted sites with or without S9.6 antibody enrichment. Total number of independent reads and R-loop footprints (peaks) are shown for each tested locus and for each strand. The percentages of R-loop peaks are indicated at right in each case.

| Gene | strand | No DRIP enrichment |  |  |  |  |  | Post DRIP |  |  |  |  |  |
| --- | --- | --- | --- | --- | --- | --- | --- | --- | --- | --- | --- | --- | --- |
|  |  | (+ ) strand |  |  | (- ) strand |  |  | (+ ) strand |  |  | (- ) strand |  |  |
|  |  | Total reads | R-loop peaks | % peaks | Total reads | R-loop peaks | % peaks | Total reads | R-loop peaks | % peaks | Total reads | R-loop peaks | % peaks |
| *CALM3* | + | 15121 | 66 | **0.4** | 1692 | 0 | 0.0 | 2729 | 351 | **12.9** | 776 | 10 | 1.3 |
| *FUS* | + | 5111 | 226 | **4.4** | 14489 | 3 | 0.0 | 2744 | 1457 | **53.1** | 3807 | 0 | 0.0 |
| *PIN4* | + | 1912 | 110 | **5.8** | 6001 | 1 | 0.0 | 926 | 590 | **63.7** | 4851 | 0 | 0.0 |
| *RPL13A* | + | 4885 | 300 | **6.1** | 5654 | 14 | 0.2 | 869 | 180 | **20.7** | 1297 | 0 | 0.0 |
| *RPS24* | + | 2373 | 258 | **10.9** | 5327 | 1 | 0.0 | 1992 | 1194 | **59.9** | 3968 | 0 | 0.0 |
| *SNRPN70* | + | 931 | 318 | **34.2** | 3070 | 19 | 0.6 | 1994 | 1007 | **50.5** | 3073 | 5 | 0.2 |
|  |  | **30333** | **1278** | **5.9** |  |  |  | **11254** | **4779** | **51.8** |  |  |  |

**Figure 2:**
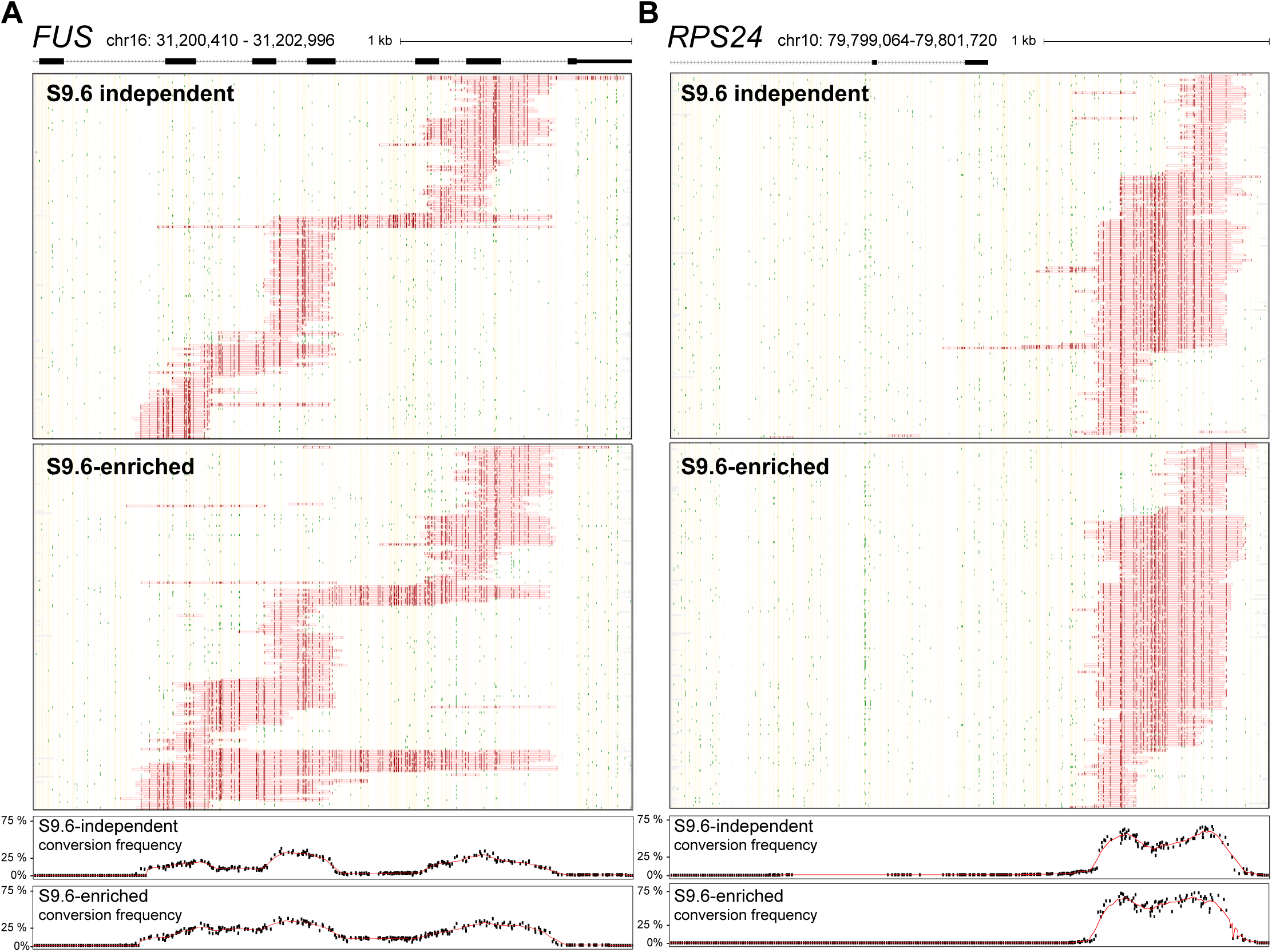
Comparison of native and DRIP-enriched SMRF-seq profiles. **a.** SMRF-seq profiles are shown for the non-template strand of the *FUS* amplicon without and with S.6 enrichment (top and bottom, respectively). 226 molecules are shown in both panels corresponding to the totality of footprints in the absence of S9.6 and to a random subsample after S9.6 enrichment. The plots below depict the aggregate C to T conversion frequencies observed for each sample. **b.** Similar data as in previous panel for the *RPS24* locus. 258 independent molecules are shown in both panels corresponding to the totality of footprints in the absence of S9.6 and to a random subsample after S9.6 enrichment.

Given that S9.6 enrichment permits a tenfold higher recovery and thus coverage, we expanded the range of loci studied using DRIP followed by SMRF-seq to a total of 24 R-loop-prone single loci (Table S1), representing a total of 78.7 kb of genomic space. This included a range of CpG island promoter, gene body, and terminal regions chosen because prior genomic mapping indicated that these regions were R-loop prone [3]. A total of 10,429 high-confidence R-loop footprints were generated, representing the most comprehensive characterization of R-loop formation with near nucleotide, strand-specific, and single-molecule resolution. As expected, SMRF-seq footprints were overall highly strand-specific and distributed on the non-template strand for transcription (Table S1). In a few instances (*KAT5, RPL4*, and *WDR3*), footprints were observed on both strands. In all three cases, the corresponding portions of these loci could be transcribed on both strands as a result of convergent or divergent transcription units, and footprints were in fact stranded with respect to transcription (Figure S5).

To determine if the ssDNA conversion footprints were caused by an RNA:DNA hybrid, we treated an S9.6-enriched population of DNA molecules with purified RNase H prior to non-denaturing sodium bisulfite treatment. RNase H treatment caused a 98.2% reduction in the proportion of footprint-carrying molecules (Table 2). The few remaining footprint-carrying molecules displayed short, mostly random footprints that are consistent with incomplete RNase H digestion. Thus, the sensitivity of the non-template DNA strand to bisulfite was dependent on the formation of an RNA:DNA hybrid necessarily involving the bisulfite-inaccessible template DNA strand. This, in turn, strongly suggests that the structures surveyed here were genuine three-stranded R-loops composed of an RNA:DNA hybrid and a looped-out DNA single-strand. Overall, SMRF-seq allows the effective characterization of single-molecule R-loop footprints at high resolution and ultra-deep coverage. In every case, R-loop footprints were co-directional with transcription; they were enriched with the S9.6 antibody and were RNase H-sensitive, as expected from the formation of genuine co-transcriptional R-loops.

**Table 2:** Summary of SMRF-seq data comparing footprinted sites with or without RNase H pretreatment. The total number of independent reads and R-loop footprints (peaks) are shown broken down by strand for each locus. The overall percentages of R-loop footprints with and without RNase H pretreatment are shown below.

| Gene | Strand | No RNase H |  |  |  | RNase H-treated |  |  |  |
| --- | --- | --- | --- | --- | --- | --- | --- | --- | --- |
|  |  | Total (+) strand reads | (+) strand peaks | Total (-) strand reads | (-) strand peaks | Total (+) strand reads | (+) strand peaks | Total (-) strand reads | (-) strand peaks |
| *CALM3* | + | 1598 | 199 | 355 | 6 | 1685 | 36 | 173 | 0 |
| *FUS* | + | 2037 | 1084 | 2488 | 0 | 1052 | 0 | 2500 | 5 |
| *SNRPN70* | + | 1450 | 767 | 2060 | 5 | 1092 | 0 | 1778 | 0 |
| *MRFAP1L1* | - | 1833 | 0 | 762 | 491 | 1249 | 0 | 1018 | 0 |
|  |  | Total stranded peaks |  | 2541 |  | Total stranded peaks |  | 36 |  |
|  |  | Total stranded reads |  | 5847 |  | Total stranded reads |  | 4847 |  |
|  |  | **% peaks** |  | **43.46%** |  | **% peaks** |  | **0.74%** |  |

### R-loops typically extend for a few hundred base pairs but can reach kilobase lengths

SMRF-seq data allowed us to precisely determine the length distribution of individual R-loops over a large collection of footprints and loci. Peak lengths varied between loci (Figure 3A) but in the majority of cases, they ranged from 100 bp, corresponding to our minimal length threshold up to 450 bp, corresponding to about three nucleosome length equivalents. When analyzed collectively, median R-loop lengths were longer for promoters (347 bp) than gene bodies (329 bp) and terminal regions (264 bp) (Figure 3B). For a few individual loci, particularly promoters, median R-loop lengths could be significantly larger. For *GADD45A*, which we analyzed almost along its entire length, the R-loop median length was 700 bp, with structures covering promoter and gene body, as predicted by DRIPc-seq (Figure S6A). Similarly, for the *PIN4* promoter, the median R-loop length was 800 bp (Figure S6B). Thus, some loci appear capable of giving rise to longer R-loop structures. For the majority of loci surveyed, a small subset of kilobase-length R-loops were recovered, contributing to a long tail in the length distribution (Figure 3). In some instances, these structures exceeded 2 kb, with the longest contiguous footprints in the dataset reaching 2.7 kb for the *PIN4* locus, equivalent to over 15 nucleosome lengths. This indicates that on occasion, R-loops can extend to great lengths and remain stable enough to be detected in our assay.

**Figure 3:**
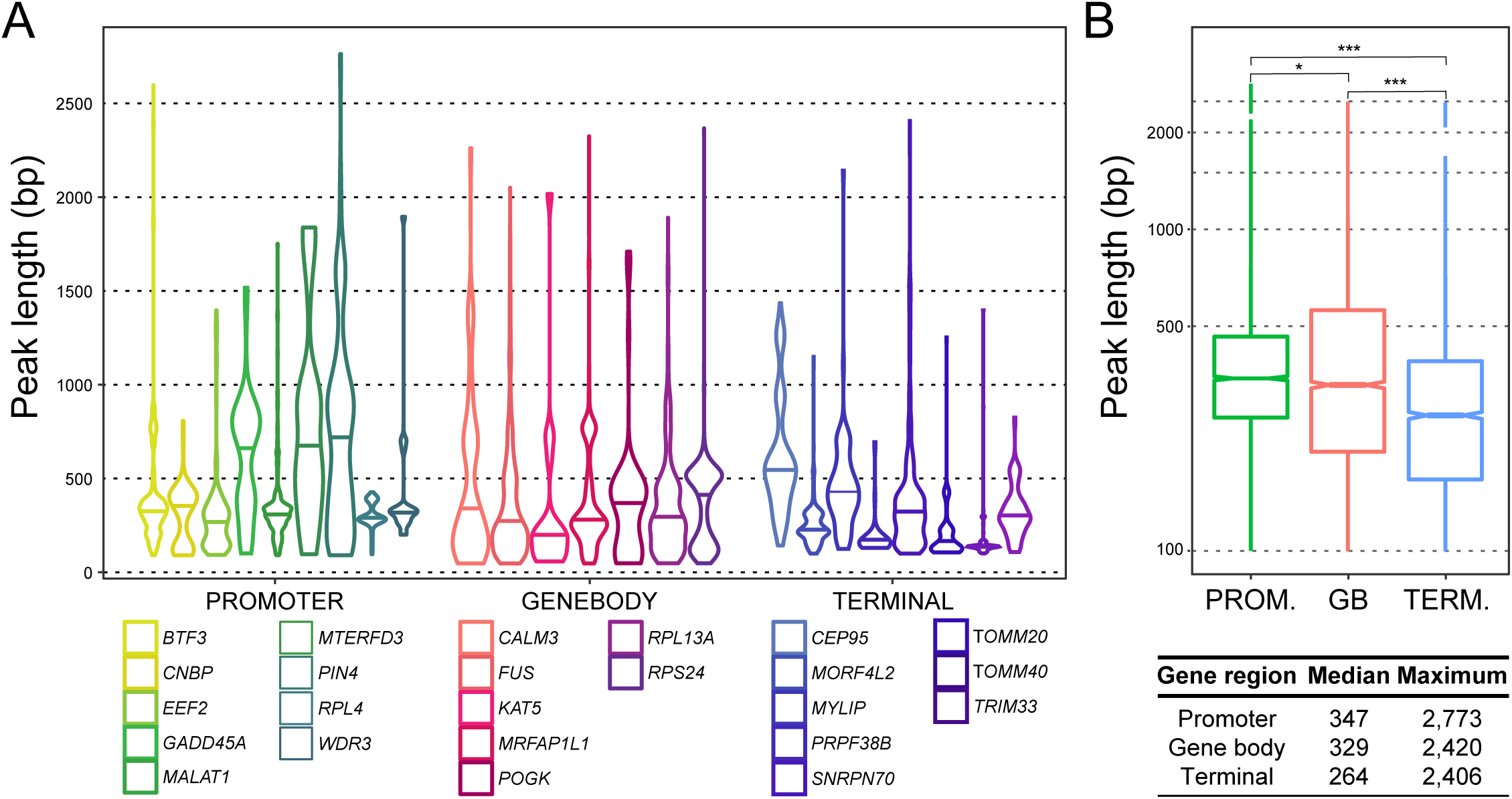
R-loop length distribution. **a.** Peak lengths are shown as violin plots for each locus. **b.** Aggregate lengths for promoter, gene body and terminal regions are displayed as box plots and the median and maximal values are indicated below.

### R-loops form overlapping molecular clusters spread over larger R-loop-prone zones

As was already evident for the *FUS* locus (Figure 1) and was confirmed for all loci (Figures 2, S2, S6), R-loops formed non-randomly over series of molecular clusters. These clusters possessed distinct starts and stops and often showed overlapping patterns of distribution. Such overlaps were caused by the existence of multiple preferred initiation sites and by a tendency for structures to extend until variable termination hotspots once initiated. The observation that R-loops aggregate to form overlapping molecular clusters provides a clear rationale for how structures of “modest” sizes can, together, create large kilobase-size R-loop-prone zones, as detected in population-average DRIPc-seq data [3]. When compared to fine gene structure annotations, R-loop footprints in most instances mapped to intronic sequences and often spread across intron-exon boundaries (Figures 1, 2, S2, S6). This argues strongly that R-loops are primarily formed on pre-mRNA before processing of the nascent transcript by splicing factors. Detailed analysis of R-loop initiation and termination regions did not reveal any correlation with splicing junctions, further suggesting that splicing does not constrain R-loop boundaries. Similarly, R-loop footprints were clearly not limited to the first exons of genes, as reported recently [27], and were readily observed over gene body and terminal regions, in accord with DRIPc-seq data. Even in instances where R-loops appeared to form predominantly in the vicinity of the first exon, most structures in fact spread into the first intron (Figure S6C,D).

It has been suggested that the time between cell lysis, DNA extraction and processing, and bisulfite probing could cause R-loop patterns to shift [27]. To test this, we modified our procedure so that bisulfite probing was conducted a mere 25 min after cell lysis without any DNA extraction or fragmentation. Loci of interest were PCR amplified immediately after non-denaturing bisulfite treatment without any S9.6 enrichment. Overall, the conversion patterns observed after direct conversion were similar to those obtained after delayed conversion (Figure S9), suggesting that R-loop distributions do not significantly shift during DNA extraction.

### RNA polymerase I-mediated R-loops form over ribosomal RNA genes

rDNA gene arrays represent the most abundantly transcribed regions of the human genome and have historically been considered a prominent source of R-loops in *E. coli*, yeast, and human cells [28-30]. To date, however, RNA Polymerase I-driven R-loops over rRNA genes have never been characterized at the single-molecule level. Here, we used SMRF-seq with and without S9.6 enrichment to characterize R-loops over a ∼2 kb amplicon covering the 18S RNA gene. As observed for RNA polymerase II-driven genes, R-loops defined a range of discrete overlapping molecular clusters over the region (Figure 4A). Thus, the formation of such clusters is observed for both RNA Pol I and RNA Pol II-driven R-loops, as well as R-loops generated *in vitro* by the bacteriophage T3 RNA polymerase [31]. 18S RNA R-loop lengths were within the range measured for protein-coding genes with evidence for kilobase-length structures (Figure 4B). A small but measurable proportion of reads (∼4%) carried two ssDNA footprints separated by a gap (Figure 4A). These events could reflect that two transcription complexes were engaged over the region, each one leading to one event of R-loop formation. This could be consistent with the high rates of transcription typically observed at rDNA regions. Alternatively, we cannot rule out that these multi-patch structures could result from the partial collapse of an original larger R-loop. The overwhelming majority of structures profiled from RNA polymerase II-driven genes only carried one R-loop per molecule (∼99%). Overall, SMRF-seq confirmed genuine R-loop formation over rRNA genes mediated by RNA polymerase I.

**Figure 4:**
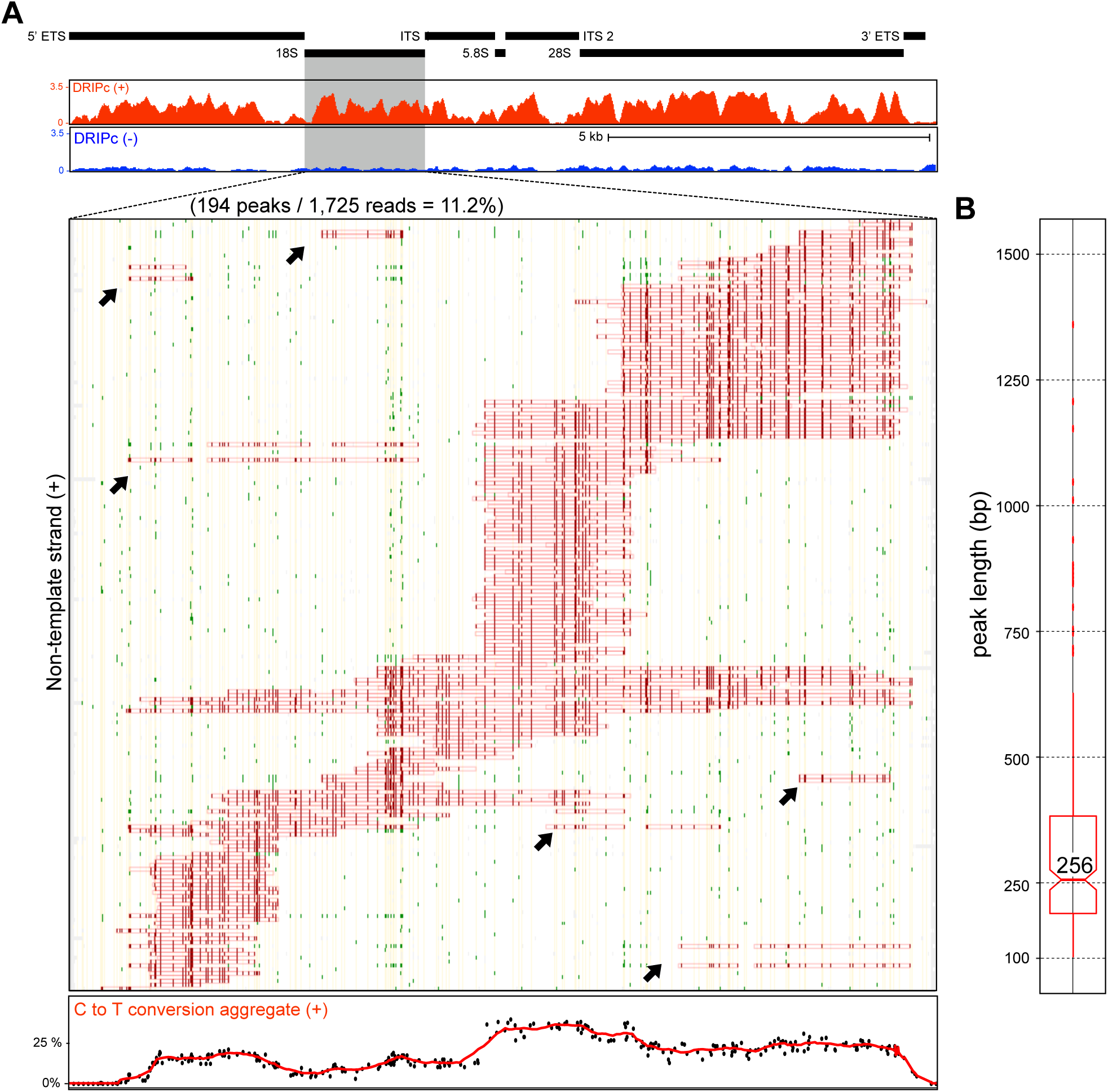
a. RNA Polymerase I-mediated R-loop formation over the 18S region. **a**. Black boxes highlight the gene arrangement over the rDNA region (top). The DRIPc-seq profile over this region is shown below for both positive and negative strands in red and blue, respectively. R-loop footprints collected with SMRF-seq over the 18S regions are shown for the non-template strand. Color codes are as described in Figure 1. The aggregate C to T conversion frequency is graphed at the bottom. Possible “double R-loops” are highlighted on panel A by black arrows. **b.** Peak length distribution for non-template strand 18S R-loops.

### Short template strand ssDNA patches are found at R-loop junctions

R-loops are bounded by a pair of junctions between the structure *per se* and the surrounding dsDNA. It is possible that short patches of ssDNA exist on the template DNA strand at these junctions but this possibility has not been examined due to the lack of deep coverage datasets. Using our peak calling algorithm, we could not identify genuine C to T conversion footprints on the template strand even when lowering the minimal length threshold to 10 bp, suggesting that these patches, if they exist, are short. Nonetheless, the identification of well-populated and well-defined R-loop clusters enabled us to ask whether a subset of molecules carried ssDNA patches around these clusters’ edges on the template strand (see Methods for details). In almost every single case analyzed, spanning numerous loci and clusters, we were able to identify a significant portion of template strand molecules carrying such junctional ssDNA patches (Figure 5). Analysis of these junctions showed that they carried from one to three highly reactive cytosines, suggesting that junctions are short. In every case, similar ssDNA patches could not be identified if the positions of the non-template strand clusters were shuffled (p<0.05), establishing that these patches were unique to the edges of actual R-loop clusters. In some instances, a peak of ssDNA reactivity was observed on the template strand upstream of the annotated start of the R-loop cluster on the non-template strand (Figure 5, top, *PIN4* locus). Such events likely reflect that the R-loop cluster in fact initiated at this junction but could not be detected on the non-template strand due to a lack of cytosines. Thus, template strand reactivity may in some instances allow a more precise annotation of R-loop boundaries.

**Figure 5:**
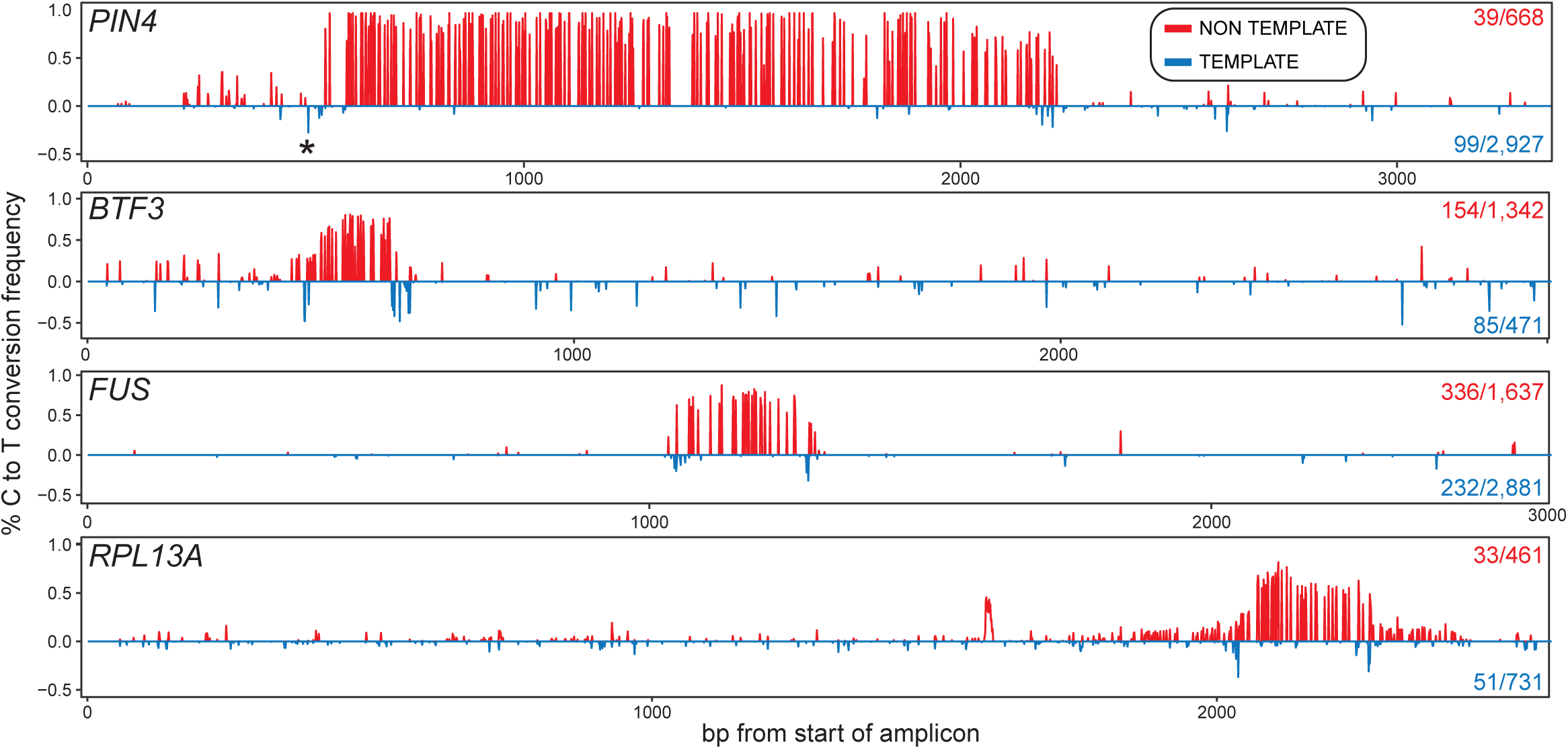
R-loop clusters are flanked by short template-strand junction “breathing”. Each panel displays the C to T conversion frequencies per residue over a specific R-loop cluster at a given locus (indicated at left). The conversion frequencies on top (red) were derived from the non-template (displaced) strand. The number of molecules that define this cluster out of the total number of converted non-template strands are indicated at right. The conversion frequencies on the bottom (blue) were derived from template strand (RNA-bound) molecules carrying properly positioned template-strand junction spikes. The number of molecules carrying such junctions out of the total converted for that strand is shown at right. Background C to T conversion frequencies measured across each amplicon were first subtracted. Patches of ssDNA, reflected in a local spike in C to T conversion, are observed on the template strand at the edges of most clusters.

### R-loop cluster boundaries can only be partially accounted for by DNA sequence transitions

Our deep single-molecule footprint collection allowed us to query whether specific sequence transitions occur at the boundaries of R-loop footprints. GC skew, purine skew, and GC content have all been implicated in R-loop formation and/or elongation [2, 21, 32] and were accordingly analyzed. An unbiased analysis of sequence motifs (k-mers) was also conducted. For this, we devised three 100 bp windows located immediately before, across or inside the upstream R-loop footprint boundary and measured these DNA sequence parameters across the entire collection of footprints (Figure 6A). Three additional 100 bp windows located inside, across and outside of the downstream R-loop footprint boundary were also used to measure sequence transitions at the end of R-loops. As shown for promoter regions, totaling 3,943 individual footprints, GC skew rose from near zero upstream of footprints to positive values inside footprints (Figure 6B). An increase in AT skew and therefore a trend towards more purine-rich RNA sequences in R-loops was also observed (Figures 6C, S7), consistent with the thermodynamic favorability of G/A-rich RNA:DNA hybrid sequences [33] and with R-loop mapping data from mammals to plants [2, 3, 10, 32, 34]. At the distal edge of footprints, reverse transitions back to the local average were not as prominent especially for GC skew. Only GC content showed a clear downward trend, attributable in this case to the fact that downstream R-loop boundaries are further away from core GC-rich CpG island promoters (Figure 6D). Promoter R-loop footprints are hence characterized by upstream trends towards higher G/A skew and reduced GC content downstream, in agreement with prior R-loop footprinting data [21].

**Figure 6:**
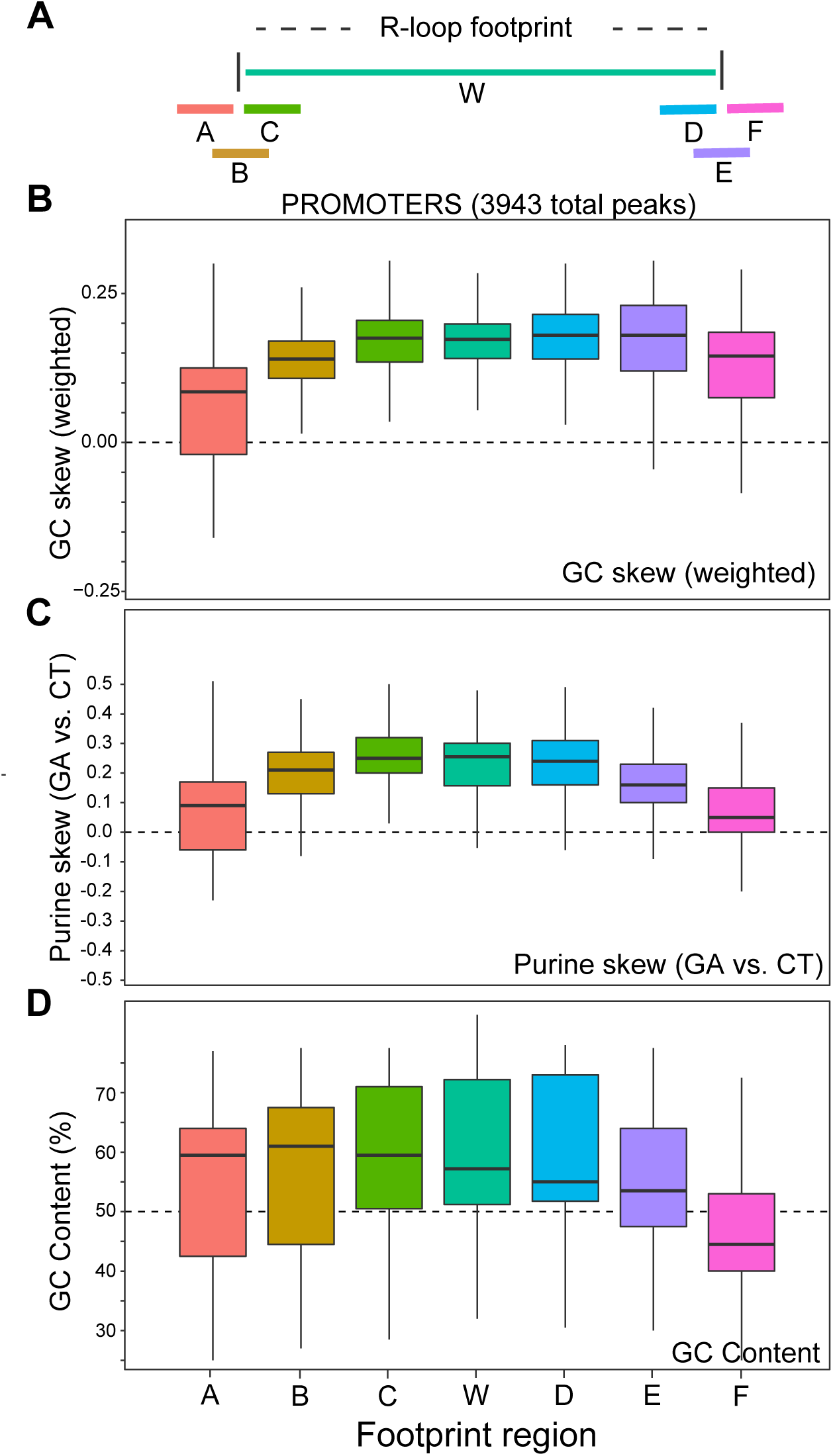
Analyzing sequence transitions at R-loop edges. **a.** Schematic of the windows used for DNA sequence analysis; each window was 100 bp long. **b-d.** Boxplots depicting variation in GC skew, purine skew and GC content for a broad range of promoter R-loop footprints across windows spanning R-loop proximal and distal ends.

Despite these overall trends, we noticed that even within the promoter dataset, some loci showed more pronounced DNA sequence boundary transitions than others. At the *EEF2* locus, strong shifts in GC skew were observed at both proximal and distal R-loop edges (Figure S7A), consistent with a major role for DNA sequence in driving structure formation. At the *PIN4* locus, however, GC skew overall was highly positive but did not show clear transitions at R-loop boundaries (Figure S7B). Thus, boundaries are not strictly defined by DNA sequence parameters. For gene body and terminal regions, GC skew, GC content and purine skew transitions were consistently observed especially at the proximal edge of R-loop footprints but the trends were weaker than those described at promoters (Figure S7C,D). Once again, some loci showed clear expected trends while others did not, resulting in high variability. In addition, no significantly overrepresented k-mer could be identified in any dataset other than a general tendency towards GA-rich sequences inside R-loop footprints. Overall, this analysis revealed that while R-loop footprints delineate broadly favorable DNA sequences, the transitions into and out of an R-loop footprint are not necessarily accompanied by systematic and predictable shifts in DNA sequence properties or with any specific motif. As an illustration, we note that advanced sequence-based R-loop prediction software such as R-loopDB [35] failed to predict some prominent R-loop forming regions detected both by DRIP-based methods and SMRF-seq (Figure S8). Our data further suggest that the extent to which a set of R-loop footprints displays expected R-loop favorable DNA sequence characteristics varies from locus to locus.

One way to understand R-loop distribution patterns builds on recent advances demonstrating the key role of the interplay between DNA sequence and negative DNA superhelicity in driving R-loop formation [31]. R-loops were proposed to form because they lower the energy of the DNA fiber via the relaxation of the negative superhelical stresses and the formation of favorable RNA:DNA hybrids. Since negative supercoiling is required for R-loops to form even when favorable sequences exist, R-loop prone zones were proposed to represent genomic regions that experience significant torsional stress. Factoring in DNA topological constraints lessens the requirements for DNA sequence to drive R-loop formation. The resulting model predicted *in vitro* R-loop positions with high accuracy [31]. To test whether this approach can predict the position of genomic R-loops, we used the R-looper software [31] to mathematically compute the probability of R-loop formation over amplicons analyzed by SMRF-seq. A negative superhelicity of −3.5%, which is compatible with that generated by transcription [36, 37], was assumed. Areas with significant R-loop probabilities were called from the probability distribution and the position of these predicted regions was compared to actual footprints in terms of direct intersect or distance. As a control, we shuffled the probability peaks 125 times independently. A similar approach was taken using predictions from R-loop DB. Both approaches performed well over promoter regions, which often possess strong favorable sequence signatures, with predictions being 2.8 to 8.8 higher (R-looper) and 2.9 to 6.1 times (R-loop DB) more likely to match an R-loop footprint than expected by chance (p<0.008) (Table 3). Predictions remained strong over gene body and terminal region using the topology-enabled R-looper tool, with odds of intersecting actual R-loop 6.8 to 15.9 times higher than at random. The sequence-based R-loop DB tool, however, performed poorly on a number of loci, lowering the predictive power. Overall, these results suggest that R-loops cannot simply be predicted using DNA sequence characteristics alone. Instead, factoring in DNA topology significantly boosted our ability to predict R-loops especially outside of promoter regions.

**Table 3:**
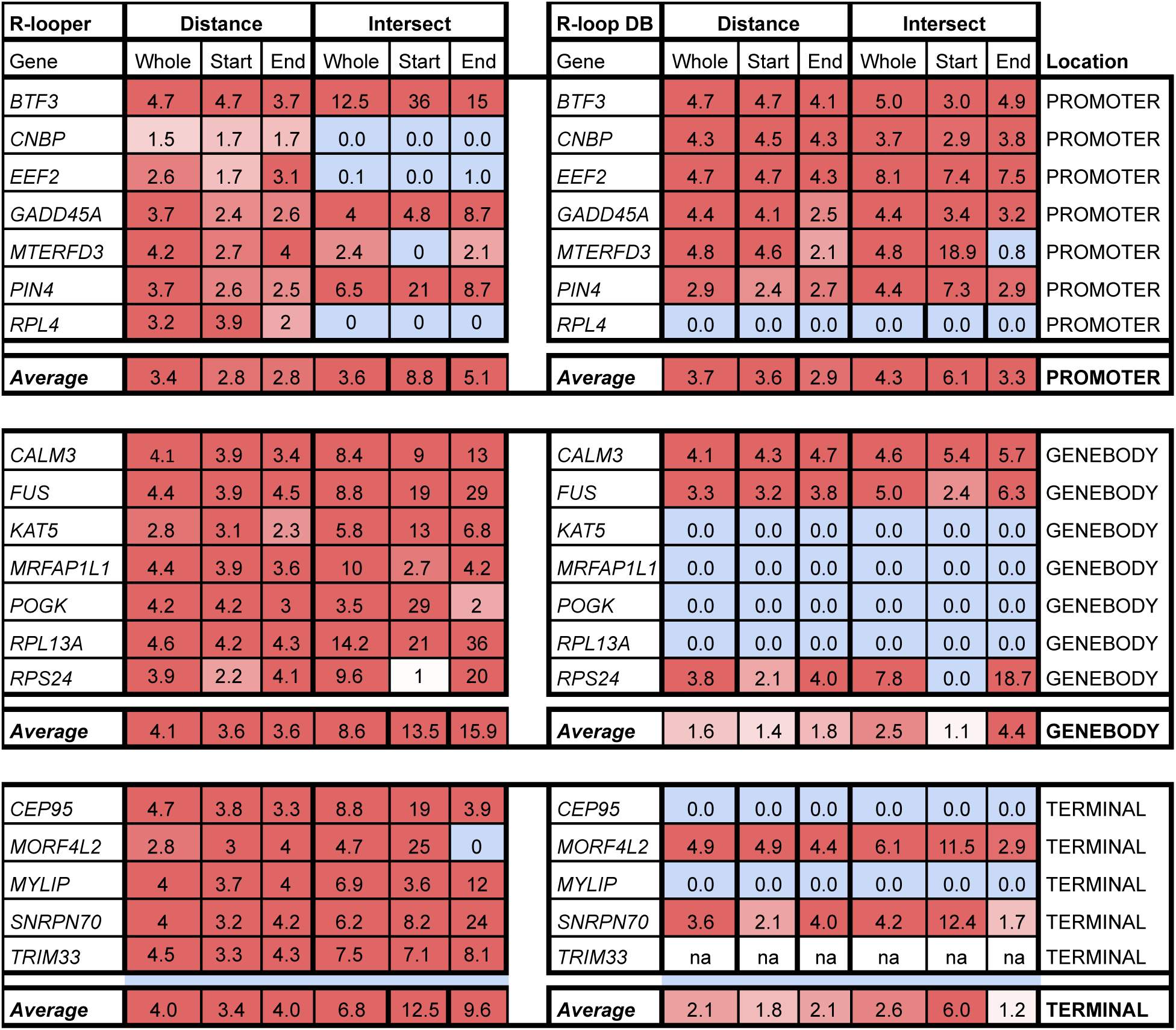
Comparison of R-loop prediction algorithms. The ability of R-looper and R-loop DB algorithms to predict R-loop clusters was analyzed for a range of footprinted loci. Values correspond to the observed over expected ratios for direct intersect or nearest distance. Expected values were generated by shuffling predictions along the amplicon 125 times. Distances or intersects were calculated either to the start (100 bp), end (100 bp) of each footprint or with the whole footprint (middle position for distance). A value of zero indicates that no match was observed or that no prediction was made. ‘na’ indicates that this region wasn’t computed to possible R-loops.

## DISCUSSION

Based on the exquisite reactivity of ssDNA to bisulfite-mediated cytosine deamination [20], SMRF-seq allows users to characterize R-loops in a strand-specific manner, at single-molecule, near nucleotide resolution and ultra-deep coverage. Importantly, SMRF-seq can be conducted independently of S9.6 enrichment, using only native PCR primers, and therefore represents an orthogonal approach to DRIP-based methods for R-loop characterization. The dataset presented here largely expands our knowledge of R-loop formation.

SMRF-seq profiles establish that three-stranded R-loop structures commonly form in the human genome. This results from two main arguments. First, non-template and template strands produce highly asymmetric bisulfite sensitivity patterns with high conversion rates only on the non-template strand for transcription. This argues that while the non-template strand is largely single-stranded and accessible, the template strand is engaged in contiguous base-pairing interactions and chemically inaccessible, save for short conversion patches located at the edges of R-loop clusters. Second, the fact that RNase H treatment abolishes the bisulfite reactivity of the non-template DNA strand indicates that the ssDNA character of that strand depends on the formation of an RNA:DNA hybrid between the template DNA strand and the nascent transcript. SMRF-seq consequently brings conclusive evidence that three strands are involved in R-loop structures, including an RNA:DNA hybrid and a largely ssDNA. The strong agreement between SMRF-seq and DRIPc-seq data at all tested loci, together with the overwhelming strand specificity, transcriptional co-directionality, and RNase H sensitivity of DRIPc-seq signal, suggests that the large majority of S9.6-based DRIP signals also correspond to three-stranded R-loop structures.

SMRF-seq individual footprints show near absolute co-directionality with transcription at all single loci and often initiate in introns and span (multiple) exon-intron boundaries. These observations are most compatible with a model whereby R-loops form upon reinvasion of the nascent transcript before binding by the spliceosome. Using bisulfite modification followed by S9.6 immunoprecipitation and sequencing (bisDRIP-seq), Dumelie *et al.*, (2017) suggested that R-loops are restricted to the first exon of highly transcribed genes and that splicing tightly controls R-loop boundaries, or alternatively, that R-loops form from spliced transcripts[27]. Similar findings could not be obtained by SMRF-seq. R-loop formation was clearly observed outside of first exons through gene bodies and over terminal regions, in agreement with bulk DRIP-based methods[3, 15, 38]. Even when R-loops were observed over the first exon of a gene, they were not limited to this region and spread downstream in the gene body (Figure S6). We suspect that the discrepancies between the two studies arose from technical issues that limited the sensitivity of the bisDRIP-seq method. In bisulfite-based approaches, the desired information derives solely from the non-template ssDNA. It is therefore essential to preserve the integrity of the three-stranded R-loop structure; even a single nick will prevent the amplification and recovery of the displaced strand. It is possible that the vigorous and lengthy shaking (2 hours at 2,000 rpm) included in the initial step of the bisDRIP procedure could have led to nicking or degradation of the displaced strand. Indeed, a harsher treatment via sonication, results in significant loss of R-loops and complete shearing of the looped-out strand, preserving only the RNA:DNA hybrid moiety ([39], Sanz *et al.*, in preparation). Consistent with this, the signal to noise ratio of bisDRIP-seq experiments was low and extracting consistent signal required the combination of numerous replicates [27]. Assuming that the displaced strand was indeed nicked or sheared during bisDRIP-seq, longer R-loops would likely be disproportionately affected, and subsequently, lost. As a result, a subpopulation of shorter, abundant, and stable R-loops would become enriched. Indeed, R-loops recovered by bisDRIP-seq were short and could be detected only for the most highly expressed genes [27]. That these R-loops tended to track with the first exon is consistent with the fact that R-loop-stabilizing GC skew matches closely with exon 1, tracking with the length of the first exon and peaking at the exon 1 / intron 1 junction [34, 40]. Overall, the agreement between two orthogonal approaches, SMRF-seq and DRIP-based methods, strongly supports the view that R-loops can form co-transcriptionally anywhere along a gene. In that view, splicing factors and other RNA binding proteins only intervene to regulate the likelihood of interaction of the nascent pre-mRNA with the DNA template, as suggested [1, 41], but not the precise pattern of R-loop formation. This finding is consistent with the recent view that intron sequences remain in nascent RNAs long after the transcription machinery transcribed them [42].

Based on the largest collection of individual R-loop footprints to date, we show that R-loops most often range from 200-500 bp in length (Figure 3), making them an order of magnitude larger than most other non-B DNA structures such as single-stranded bubbles, B/Z transitions, or triplexes that all typically range from 20-50 bp [43]. For most loci, the accretion of individual R-loops defined discrete clusters. These clusters themselves often overlapped and spread through larger, kilobase-size, R-loop-prone zones. For some loci, R-loop footprints coincided with clear R-loop favorable sequence characteristics while for other loci, they did not, suggesting that R-loops cannot be simply captured by stereotypical sequence transitions. Patterns of R-loop formation can instead be better understood when one considers DNA topology as a driving factor for R-loops [31]. If a region experiences low to mild supercoiling stress, R-loop favorable DNA sequence characteristics will play a leading role in allowing R-loops to form. This situation most likely applies to promoter regions. These loci carry conserved patterns of R-loop-favorable sequence characteristics [40] and experience only limited transcription-driven superhelical stress given their location early in the transcription unit. By contrast, if a region experiences moderate to high topological stress, the role of DNA sequence favorability as a driver of R-loop stability will be reduced in favor of DNA relaxation [31, 44]. Given that transcription-driven topological stress builds up with the length of DNA transcribed [45], the contribution of DNA topology to R-loop formation may be higher in gene body and terminal regions. This is consistent with reports that R-loop propensity correlates with gene length and gene expression [3] and that loss of topoisomerase I results in R-loop accumulation in long, highly expressed, genes for which superhelical stress dissipation is constrained [30]. We therefore suggest that loci for which prominent R-loop hotspots form over regions with moderately or poorly favorable sequence-characteristics, such as gene body or terminal gene regions, may be driven primarily by the need to relax topological stresses. Importantly, these observations imply that strictly DNA sequence-based approaches are unlikely to support accurate and robust R-loop predictions. DNA topology-enabled approaches are instead required and early indications support that these approaches perform better for gene body and terminal regions (Table 3). Accurate, global R-loop prediction will require an intimate knowledge of the intensity and distribution of DNA topology.

The notion that both DNA topology and DNA sequence cooperate to control R-loop stability has further implications. As discussed above, the relative importance of DNA sequence versus DNA topology in allowing R-loops to form can be indirectly determined by analyzing the sequence boundaries of R-loop clusters. Sharp reciprocal transitions towards R-loop-favorable DNA sequences are indicative of a preferential role for DNA sequence, while a lack of favorable sequence transitions suggest that topology dominates in driving structure formation. In addition to regulating R-loop positions, DNA topology was also shown to regulate R-loop extension, and therefore lengths [31]. This can be understood by the simple fact that once initiated, R-loops are likely to extend until the topological tension necessary to support their formation has been relaxed. Alternatively, R-loops may terminate if the underlying DNA sequence becomes highly unfavorable. Discerning between these two possibilities may be possible by comparing sequence transitions at the proximal and distal edges of individual R-loop clusters. We and others[23] have observed that distal DNA sequence features are often less well-defined than those at proximal edges and that R-loops with the same initiation site can end at multiple downstream locations. Prior observations further showed that favorable G stretches are often necessary at the proximal edges of R-loops while the structures can then extend through areas that would otherwise not support initiation [46]. These results are expected if DNA topology plays a role in regulating R-loop extension. It therefore follows that the length distribution of R-loops may inform on the level of superhelical stress that existed prior to their formation. R-loops of median length 300 bp will be able to relax at least thirty negative supercoils, which is already considerable and will cause relaxation kilobases away from the site of R-loop formation. Rare kilobase-size R-loops are expected to relax vast amounts of negative superhelicity and to have long-range effects on the topology of the DNA fiber. Overall, careful consideration of the respective roles of DNA sequence and topology allow us to propose a framework for understanding the principles of R-loop formation and for recapitulating patterns of R-loop formation at the single molecule level. For regions that exist under low to moderate superhelical stress, R-loops will concentrate over clusters defined primarily by the favorability of their initiation region. Repeated events of initiation at a given favorable hotspot will naturally lead to the formation of individual clusters. Variation in the extent of superhelical stress that exists at the time an R-loop initiates will lead to heterogeneity in R-loop extension and thus result in series of overlapping clusters, as observed. The selection of additional initiation sites throughout a larger negatively supercoiled zone will lead to further overlaps between clusters and define broader R-loop-prone zones.

## DECLARATIONS

### Availability of data and materials

SMRF-seq and DRIPc-seq datasets have been uploaded to the NCBI GEO databased under accession number GSE130726. The Gargamel computational pipeline for read mapping, strand assignment, peak calling, clustering and data visualization is available from GitHub (https://github.com/srhartono/footLoop).

### Author Contributions

MM performed SMRF-seq and analyzed the data; MM and JG assisted in the development of the computational pipeline, which was primarily developed by SRH. JG performed control experiments. LAS performed accompanying DRIPc-seq. FC devised the study, supervised progress, and wrote the manuscript with input from all authors.

### Ethics approval and consent to participate

Not applicable.

### Consent for publication

Not applicable.

### Competing Interests

The authors declare no competing interests.

#### Acknowledgments

We thank members of the Chedin lab for critical feedback and useful discussions. We also thank Tonia Brown for editing this work.

