## Supplemental Figures and Tables for "High-Throughput Single-Molecule R-loop Footprinting Reveals Principles of R-loop Formation"

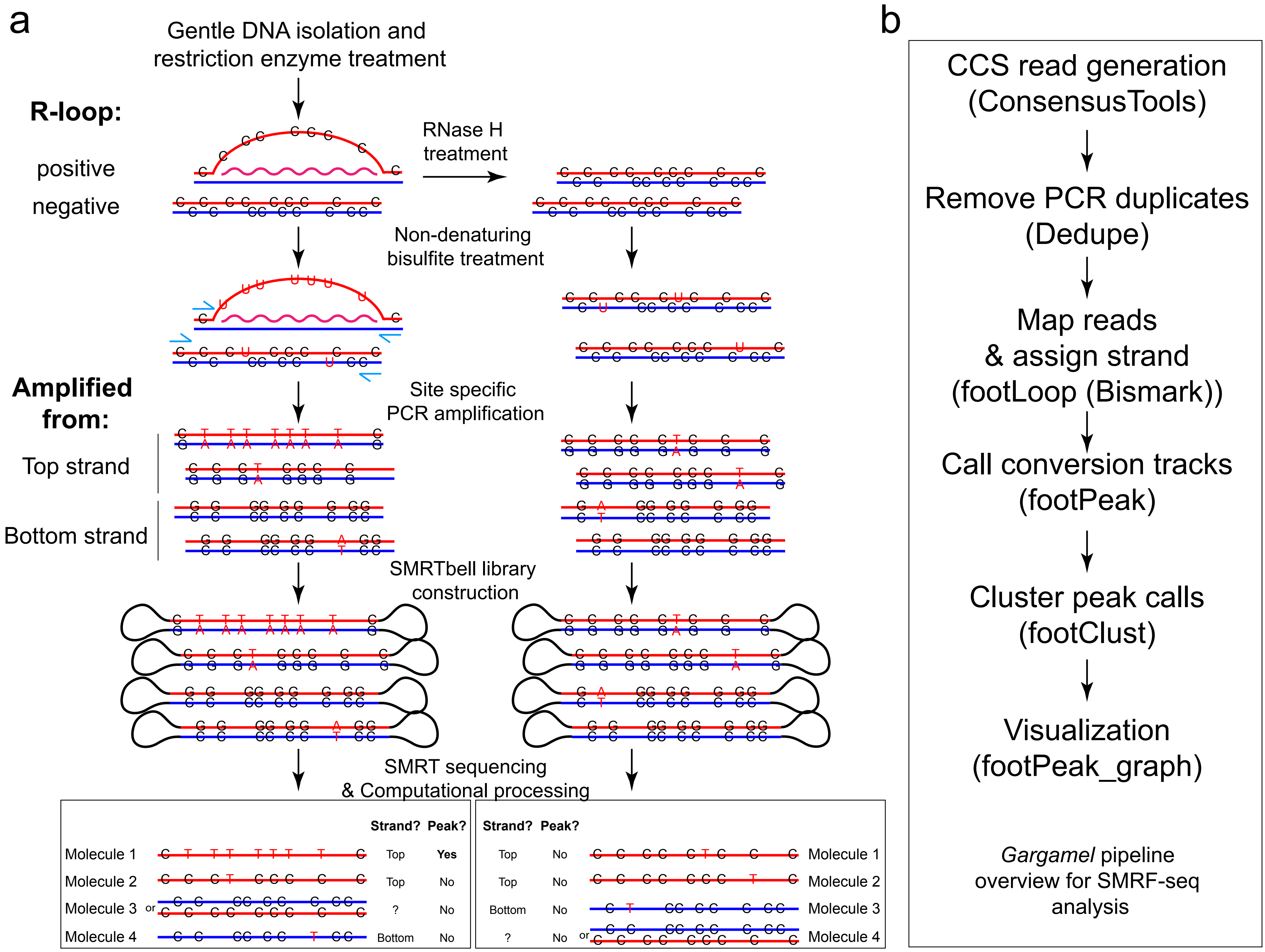


**Figure S1: a.** Overview of the experimental R-loop footprinting workflow. **b.** Overview of the computational pipeline for data analysis and visualization.

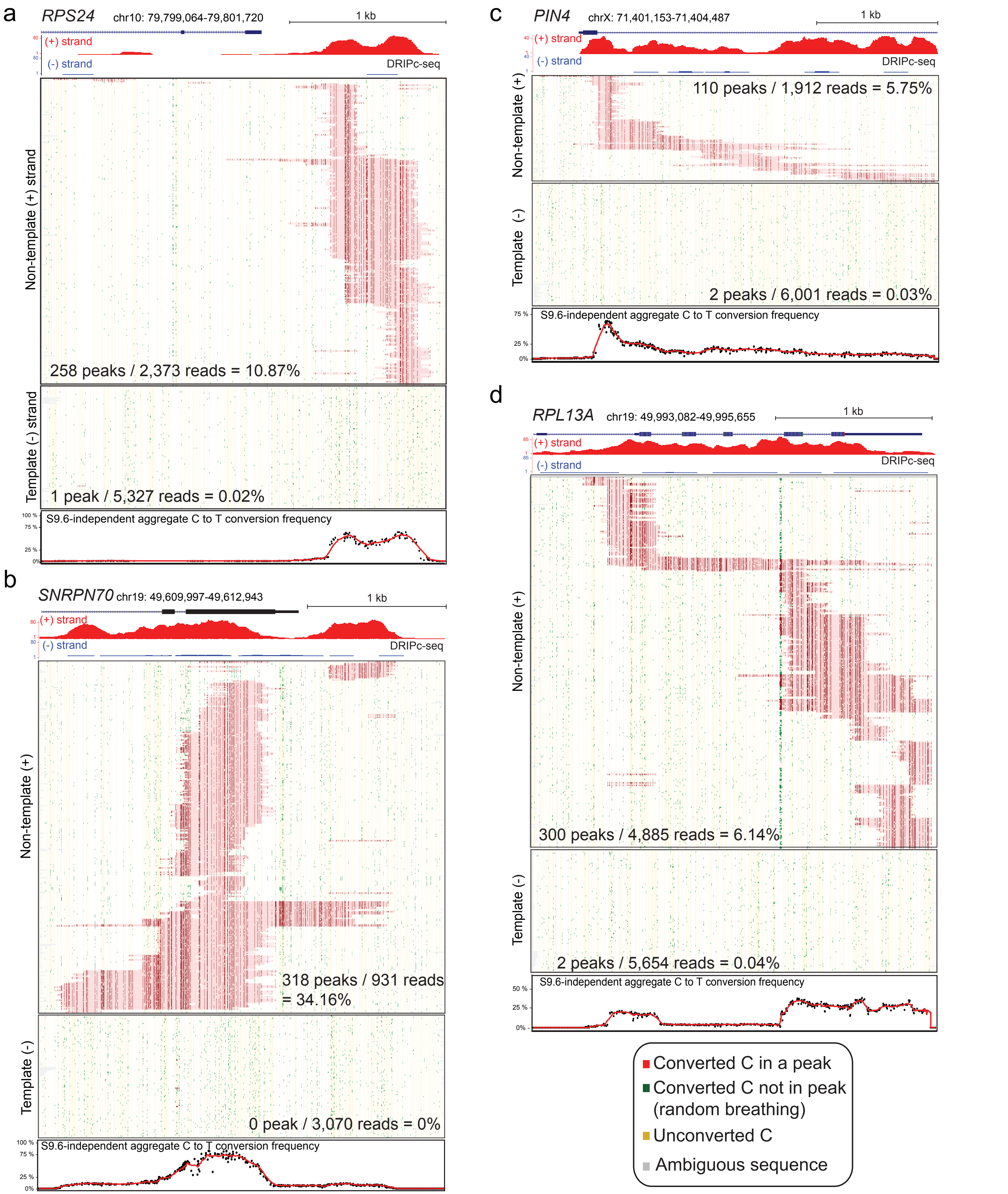


**Figure S2:** S9.6-independent R-loop footprints at the indicated loci. For each panel, the name of the locus along with the coordinates of the amplicon (hg19) are provided. The DRIPc-seq data for that region is shown at top for the (+) and (-) strands in red and blue, respectively. The R-loop footprinting data for the non-template strand and 100 reads randomly selected for the template strand are shown. The color code is shown at the bottom right of the figure. The number of R-loop footprints detected on each strand out of the total number of reads is indicated within each heatmap. The aggregate C to T conversion frequency observed for R-loops formed on the non-template strand is shown at the bottom.

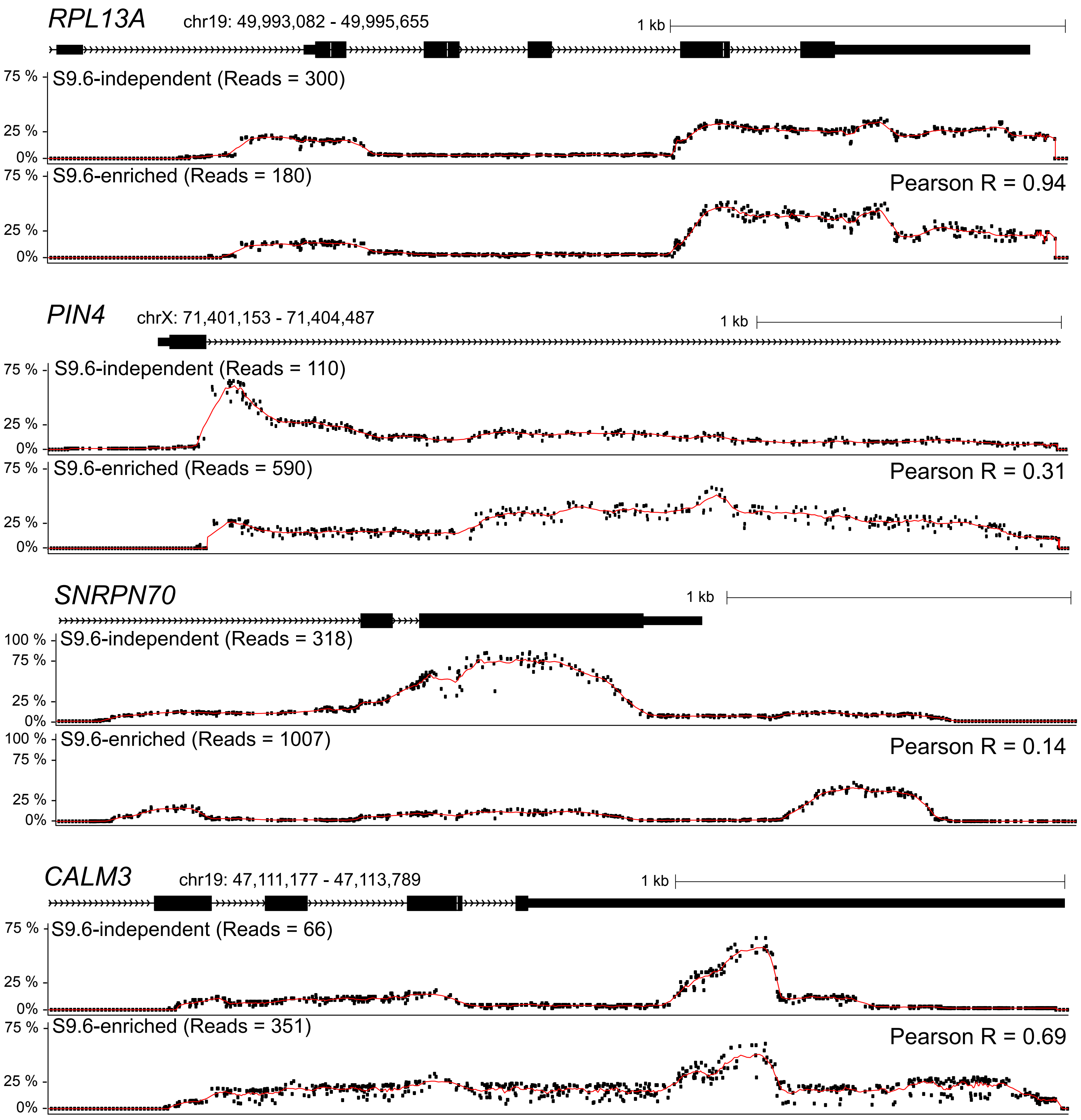


**Figure S3:** Aggregate C to T conversion frequencies with and without S9.6 enrichment are shown for the corresponding loci. Pearson correlation coefficients between both methods are indicated.

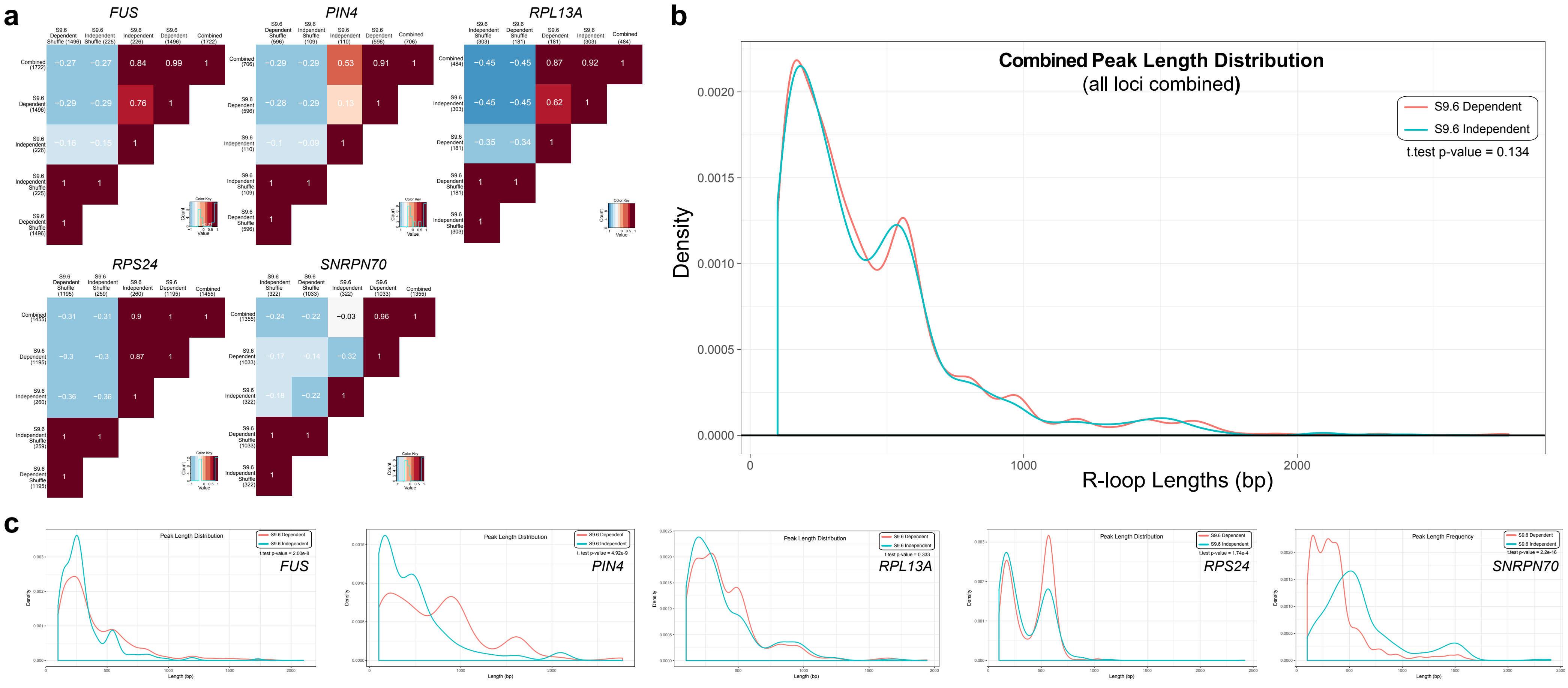


**Figure S4: a.** Positional comparison between with or without S9.6-enrichement measured by Pearson correlation coefficients. For each locus, the distribution of R-loop footprints was measured across replicates as indicated in Materials and Methods and is represented as a heatmap of Pearson correlation coefficients. The number of footprints considered in replicates is indicated in parentheses along with indication of whether the footprints were obtained with or without S9.6 enrichment. Negative controls generated by randomly shuffling the position of R-loop footprints are shown. **b.** Peak length distribution frequency histogram comparing the lengths of footprints observed with (blue line) and without (red line) S9.6-enrichement (all loci combined). Two-tailed t-test (p-value = 0.134) shows no significant difference in peak length. **c.** Peak length distribution frequency comparing the lengths of footprints with or without S9.6-enrichement at individual loci. Two-tailed t-test is shown for each.

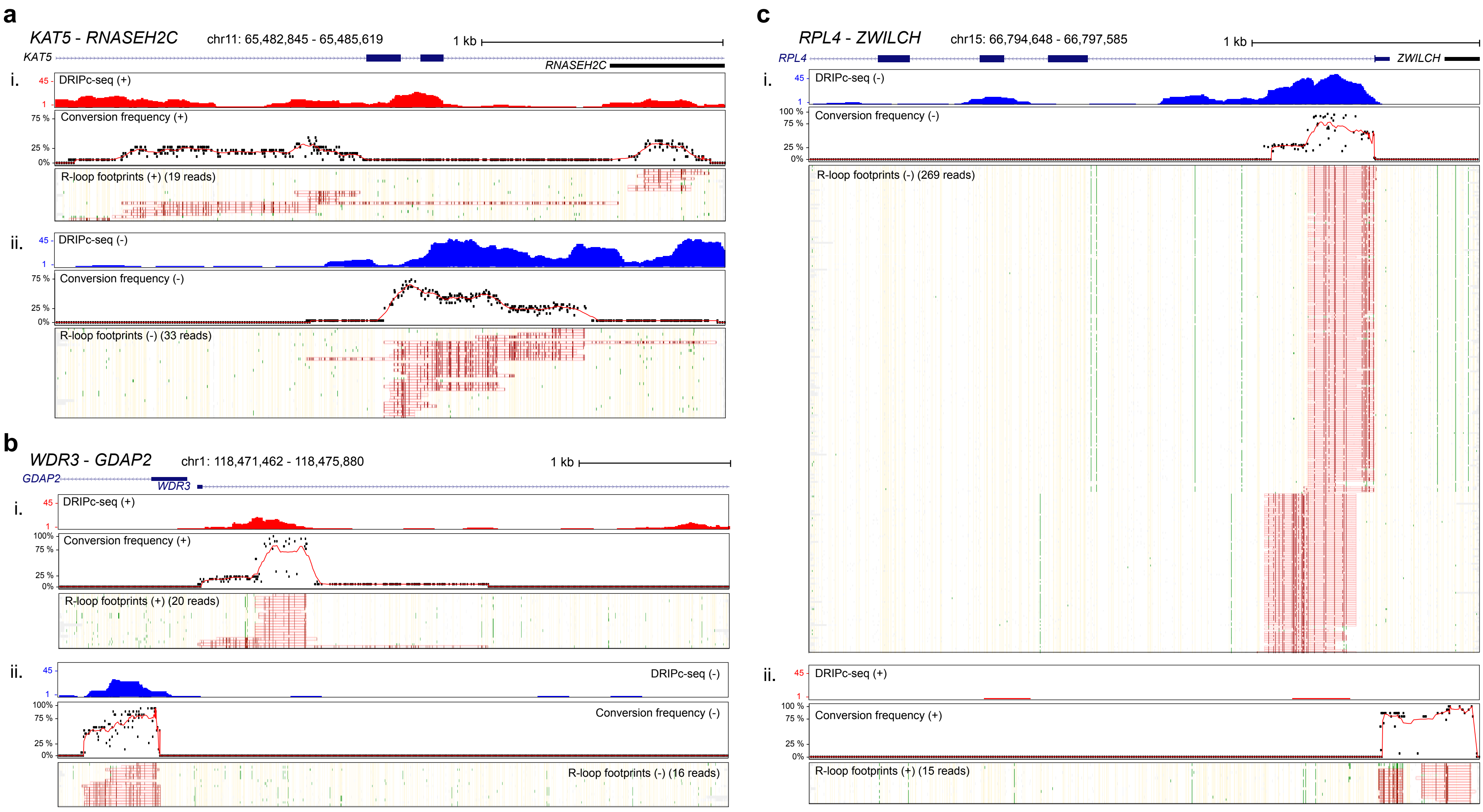


**Figure S5:** Strand-specific footprint profiles shown for divergently transcribed genes. **a.** R-loop profiles for positively transcribed KAT5 and negatively transcribed RNASEH2C. **b.** R-loop profiles for positively transcribed WDR3 and negatively transcribed GDAP2. **c.** R-loop profiles for positively transcribed ZWILCH and negatively transcribed RPL4.

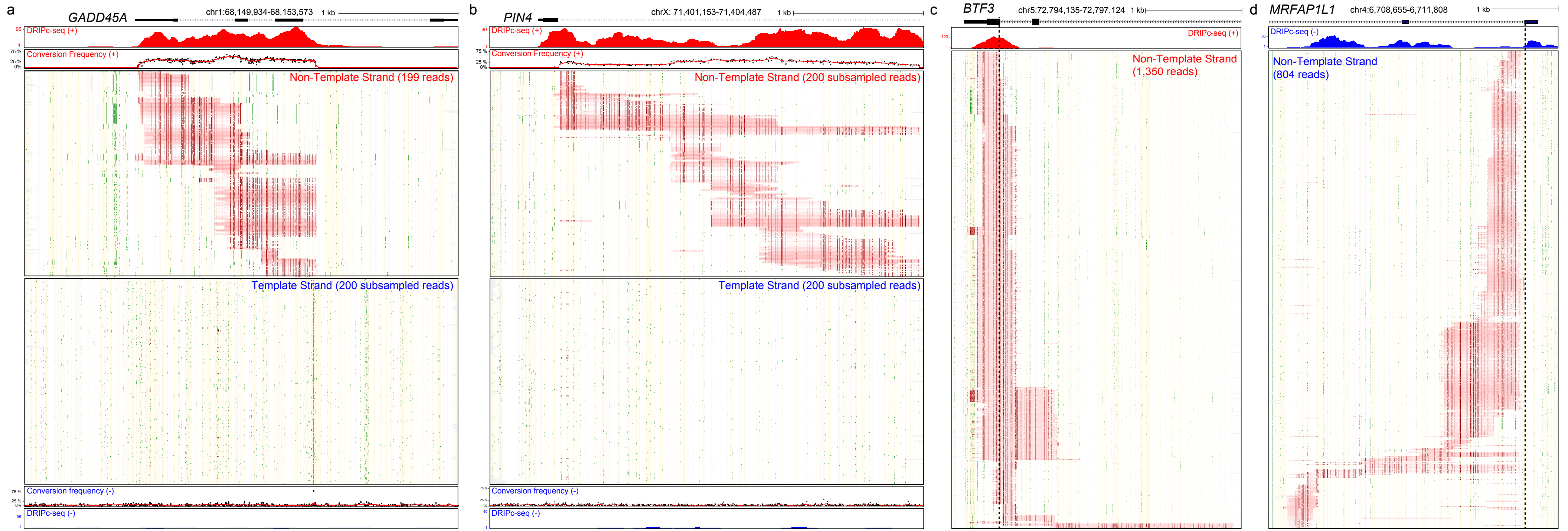


**Figure S6: a-b.** Long, contiguous R-loop footprint profiles for GADD45A and PIN4. For each panel, the corresponding gene structure, DRIPc-seq data, C to T conversion frequency, and R-loop mapping are shown for both positive (red) and negative (strand). **a.** The GADD45A gene amplicon spans 3,640 bp **b.** The PIN4 gene amplicon spans 3,335 bp. **c-d.** R-loops are not constrained to first exons. Even when R-loops tend to be observed in proximity of the first exon of a gene (panel c, BTF3; panel d, MRFAP1L1), they often spread beyond this boundary (dashed line) into the first intron. Similar results are visible for GADD45A and PIN4.

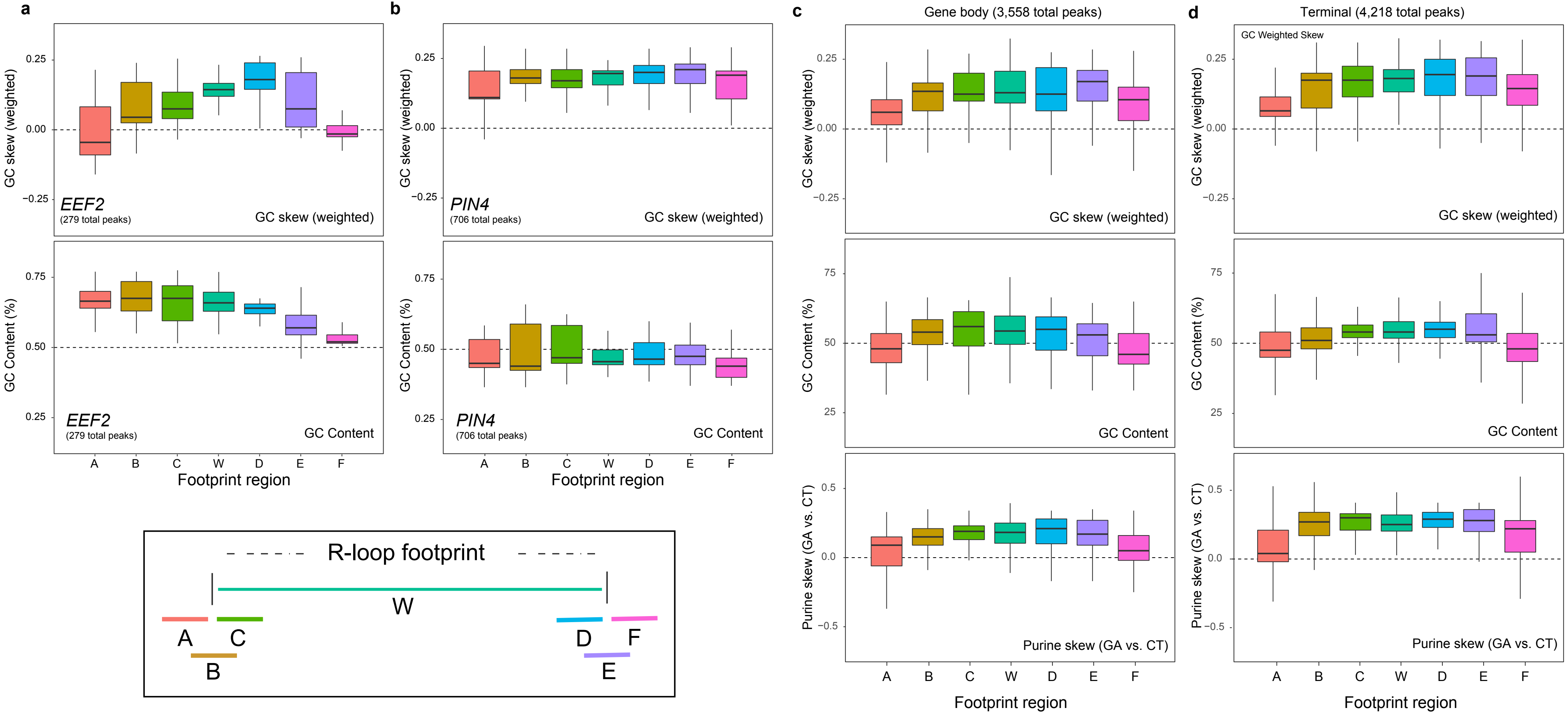


**Figure S7:** Sequence content analysis box plots for weighted GC skew, GC content, and purine skew calculated over different windows flanking R-loop proximal and distal edges. **a.** Sequence profile for EEF2 showing weighted GC skew (top) and GC content (bottom). **b.** Similar profile shown from previous panel for PIN4. **c.** Sequence profile for all gene body R-loop forming loci. **d.** Same as previous panel for all terminal R-loop forming loci.

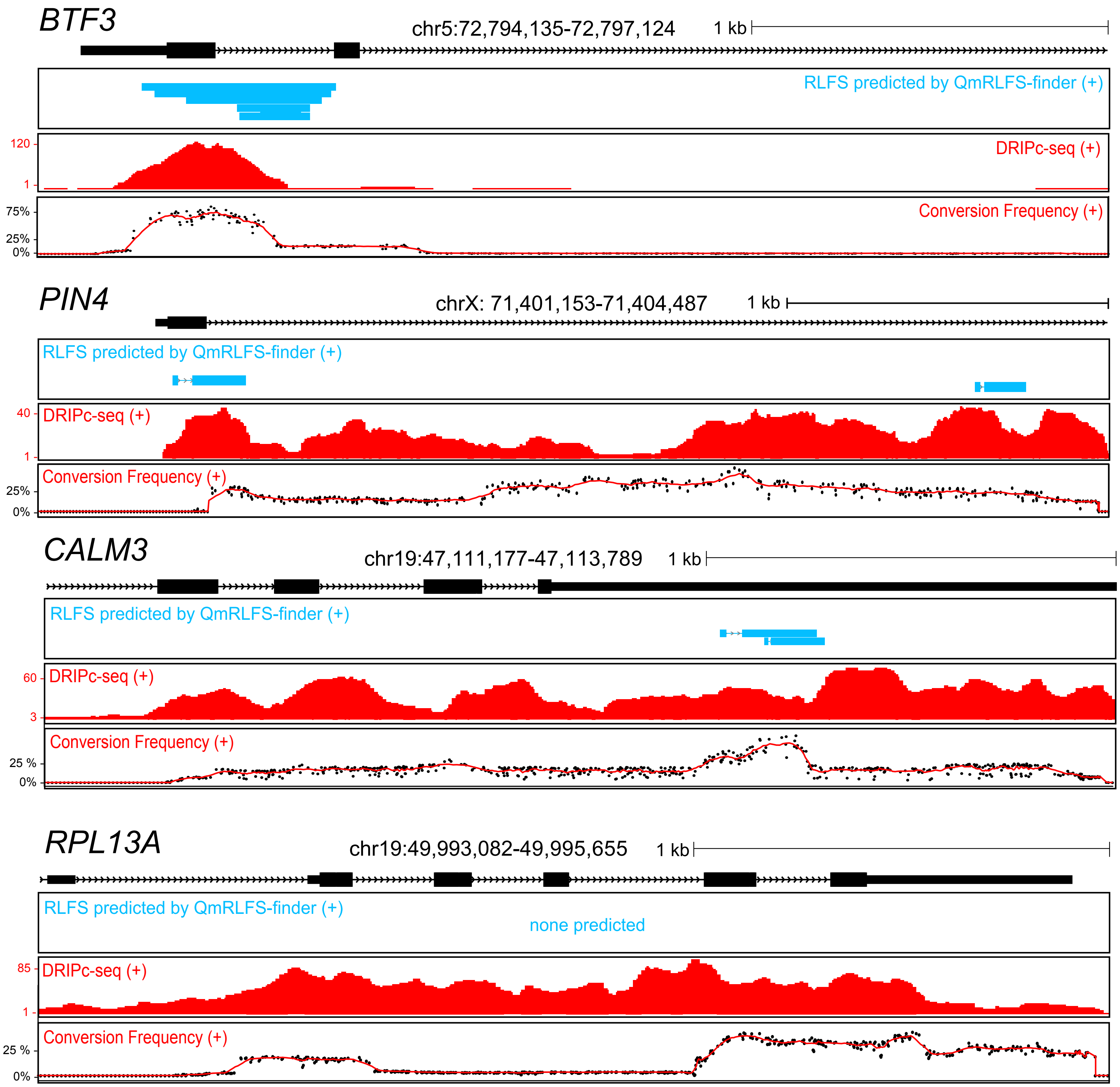


**Figure S8:** Comparison between DNA sequence-based R-loop prediction from R-loopDB (top graph) and observed R-loops from DRIPc-seq (middle graph) and SMRF-seq (bottom graph) at the indicated loci. While R-loop formation at the BTF3 promoter was correctly predicted, other loci show to variable extents that a strictly DNA sequence-based approach is insufficient to predict strong sites of R-loop formation.

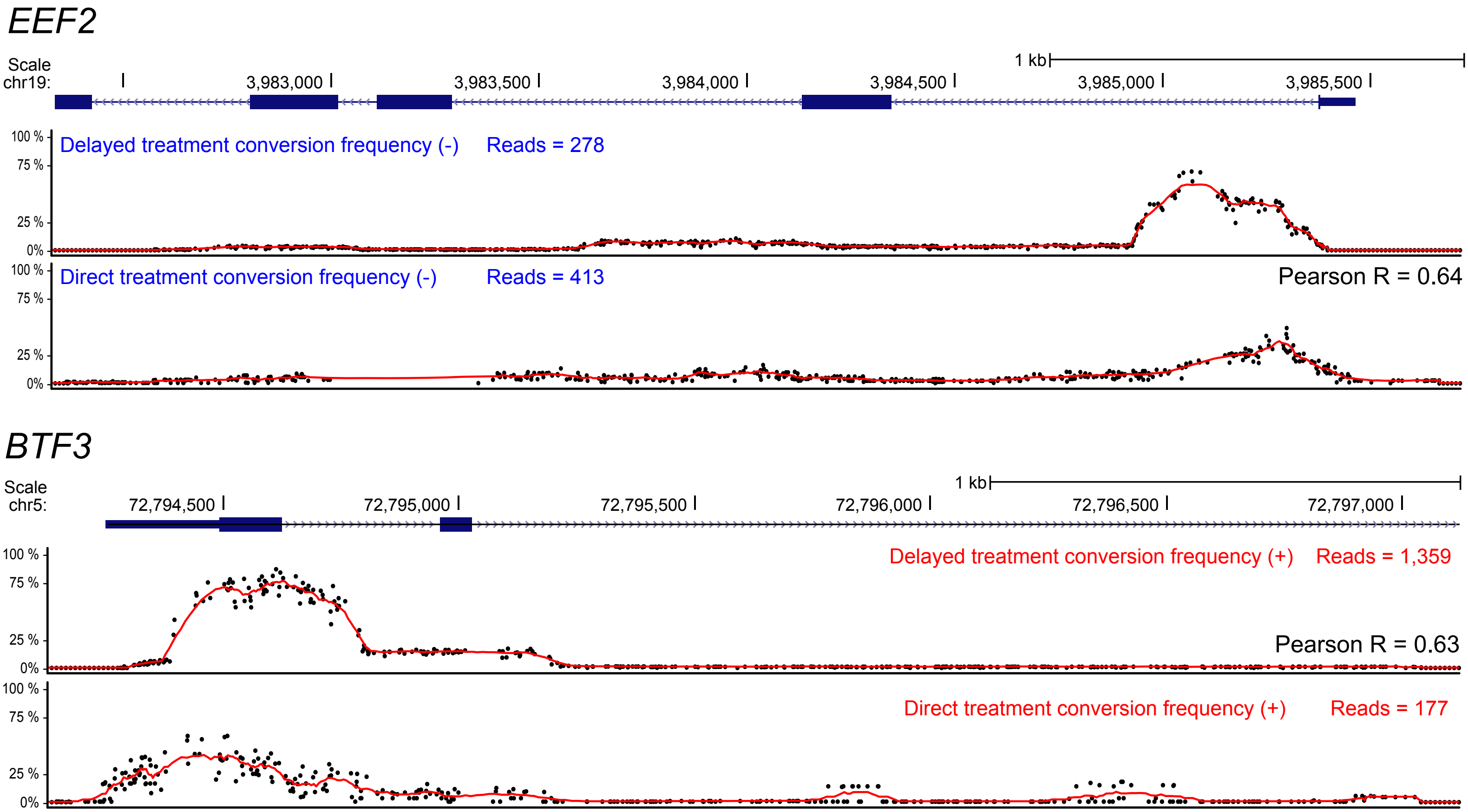


**Figure S9:** Direct bisulfite conversion. Comparison of C to T conversion frequencies between delayed and direct bisulfite conversion treatment for the EEF2 and BTF3 loci. For each locus, the genomic region along with C to T conversion frequencies for delayed (middle), and direct (bottom) bisulfite conversion are shown along with Pearson correlation coefficients between delayed and direct treatments.

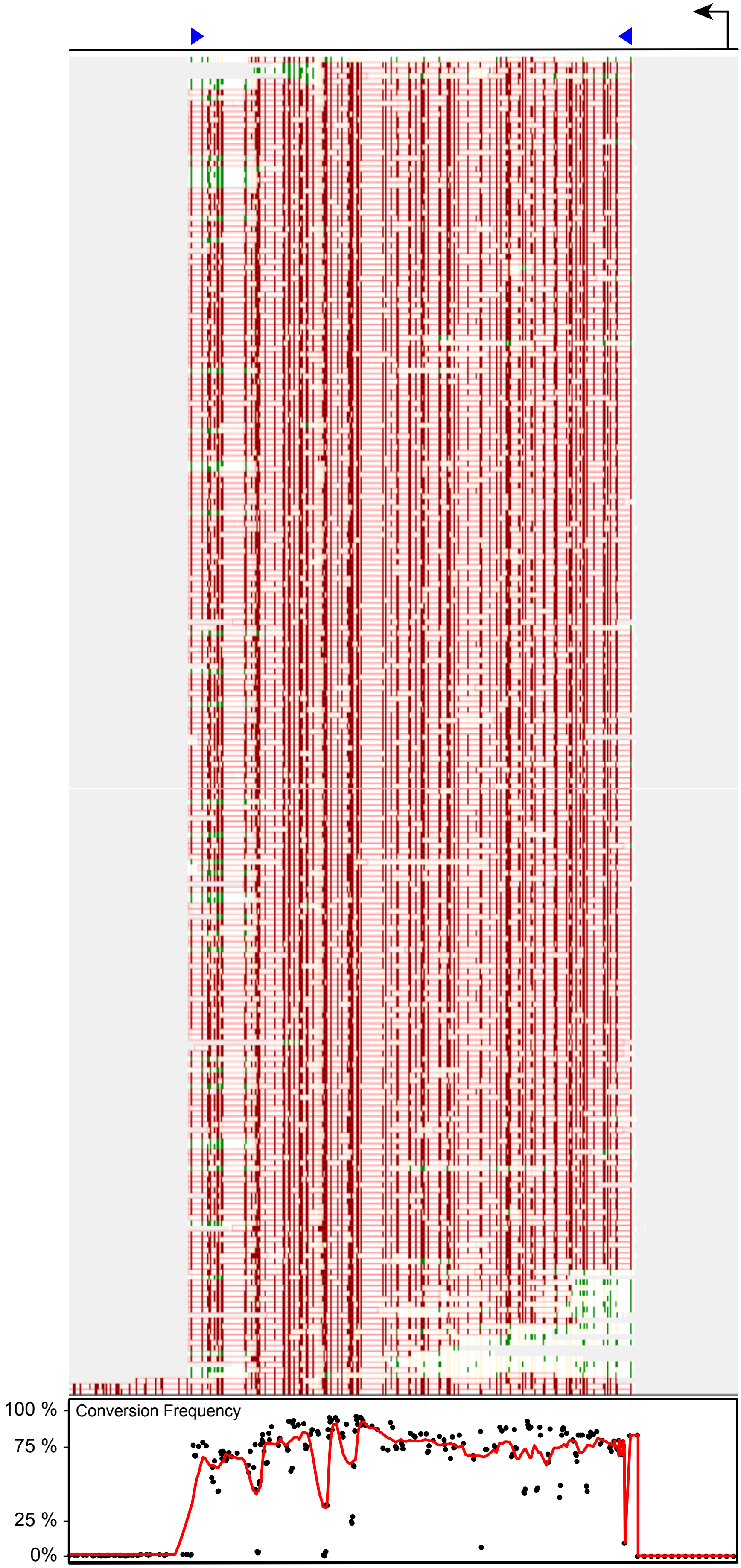


**Figure S10:** Footprints and C to T conversion profile for denatured plasmid control spike-in (n = 244). Blue arrows indicate amplicon boundaries.

| | | | | | | | (+) strand | | | (-) strand | | |
|---|---|---|---|---|---|---|---|---|---|---|---|---|
| Chrom | Start | End | Amplicon size (bp) | Gene | Strand | Locus type | Total reads | R-loop peaks | % peaks | Total reads | R-loop peaks | % peaks |
| chr4 | 6708655 | 6711808 | 3,153 | *MRFAP1L1* | - | Gene body | 2698 | 0 | 0.0 | 1224 | **803** | **29.8** |
| chr3 | 128900661 | 128903317 | 2,656 | *CNBP* | - | Promoter | 1181 | 0 | 0.0 | 848 | **176** | **14.9** |
| chr19 | 3982335 | 3985724 | 3,389 | *EEF2* | - | Promoter | 1558 | 1 | 0.1 | 1156 | **278** | **17.8** |
| chr12 | 107376879 | 107381257 | 4,378 | *MTERFD3* | - | Promoter | 1149 | 0 | 0.0 | 230 | **63** | **5.5** |
| chr15 | 66794648 | 66797585 | 2,937 | *RPL4* | - | Promoter | 3944 | 15 | 0.4 | 413 | **269** | **6.8** |
| chrX | 102929428 | 102932836 | 3,408 | *MORF4L2* | - | Terminal | 2812 | 0 | 0.0 | 1388 | **695** | **24.7** |
| chr1 | 235272022 | 235275548 | 3,526 | *TOMM20* | - | Terminal | 2703 | 0 | 0.0 | 917 | **178** | **6.6** |
| chr1 | 114931599 | 114934994 | 3,395 | *TRIM33* | - | Terminal | 583 | 0 | 0.0 | 368 | **137** | **23.5** |
| chr19 | 47111177 | 47113789 | 2,612 | *CALM3* | + | Gene body | 2729 | **351** | **12.9** | 776 | 0 | 0 |
| chr16 | 31200410 | 31202996 | 2,586 | *FUS* | + | Gene body | 2744 | **1457** | **53.1** | 3807 | 12 | 0 |
| chr11 | 65482845 | 65485619 | 2,774 | *KAT5* | + | Gene body | 1109 | **19** | **1.7** | 1041 | 33 | 3 |
| chr1 | 166818107 | 166821100 | 2,993 | *POGK* | + | Gene body | 608 | **100** | **16.4** | 1222 | 0 | 0 |
| chr19 | 49993082 | 49995655 | 2,573 | *RPL13A* | + | Gene body | 869 | **180** | **20.7** | 1297 | 0 | 0 |
| chr5 | 72794135 | 72797124 | 2,989 | *BTF3* | + | Promoter | 2007 | **1359** | **67.7** | 3440 | 0 | 0 |
| chr1 | 68149934 | 68153573 | 3,639 | *GADD45A* | + | Promoter | 547 | **199** | **36.4** | 232 | 0 | 0 |
| chr11 | 65264805 | 65268402 | 3,597 | *MALAT1* | + | Promoter | 1117 | **856** | **76.6** | 1035 | 0 | 0 |
| chrX | 71401153 | 71404487 | 3,334 | *PIN4* | + | Promoter | 926 | **590** | **63.7** | 4851 | 7 | 0 |
| chr1 | 118471462 | 118475880 | 4,418 | *WDR3* | + | Promoter | 1860 | **16** | **1** | 2432 | 20 | 1 |
| chr17 | 62531459 | 62535650 | 4,191 | *CEP95* | + | Terminal | 554 | **111** | **20.0** | 1298 | 0 | 0 |
| chr6 | 16147958 | 16151697 | 3,739 | *MYLIP* | + | Terminal | 162 | **87** | **53.7** | 420 | 0 | 0 |
| chr1 | 109242027 | 109245458 | 3,431 | *PRPF38B* | + | Terminal | 810 | **81** | **10.0** | 1658 | 14 | 1 |
| chr10 | 79799064 | 79801720 | 2,656 | *RPS24* | + | Terminal | 1992 | **1194** | **59.9** | 3968 | 8 | 0 |
| chr19 | 49609997 | 49612943 | 2,946 | *SNRPN70* | + | Terminal | 1994 | **1007** | **50.5** | 3073 | 6 | 0 |
| chr19 | 45405433 | 45408838 | 3,405 | *TOMM40* | + | Terminal | 462 | **107** | **23.2** | 582 | 0 | 0 |

**Table S1:** List of all footprinted loci with genome coordinates (hg19), locus name, strand, locus type, and summary of overall SMRF-seq data including DRIP-enriched.

| Gene | Gene Region | Forward Primer (5'->3') | Reverse Primer (5'->3') | RE mix |
|---|---|---|---|---|
| *BTF3* | Promoter | AGCAGGAGGAGGAGAATAGGG | GTCTATCACACCCAACCACCTC | N12 |
| *CALM3* | Gene body | GTCAGACCATTTTCCCCACTGT | GGAAAAGCCACTTGATCGACCT | S1 or N1 |
| *CEP95* | Terminal | ACACTGATGAAGGGAGGACAAG | AGTTCCCACCAATCACCAGA | N11 |
| *CNBP* | Promoter | CAAGAGATGAAATCGGAAAAACTT | ACCAGGCACTTTCATCTTGG | S2 or N3 |
| *EEF2* | Promoter | CCACATACATCTCGGCAAACT | ATCGTTGCTATGGTTCTCGTTC | N10 |
| *FUS* | Gene body | TTTCACATTTGCATTTTCTCTGTT | GAAAACTCTCTACCTTCCTGATCG | S1 or N6 |
| *GADD45A* | Promoter | CTCTCTGTGGAAGGTAACGACA | TTGGCATCAGTTTCTGTAATCCT | N11 |
| *KAT5* | Gene body | TAGGTGATGAAGATGCTGTG | GAGTGTCACCTGCTGTAAGA | S1 or N6 |
| *MALAT1* | Promoter | AAAGGAAGACCTAGACTGAAAATGG | TAAGCCTGAAAAAGAGAAACCTACA | N16 |
| *MORF4L2* | Terminal | CATTTGTCCTCCTCCCACTTAC | CTCCTCCCTCACCAATCTTTTT | N8 |
| *MRFAP1L1* | Gene body | CCAAAACGTGTATCTGAGCTTG | AGCTTTAGATGAAGACGCGAAC | S2 or N4 |
| *MTERFD3* | Promoter | GCTCATTTTCAGTTGTTGATGG | ATTATTTTGGGAACAGTGATTGG | N8 |
| *MYLIP* | Terminal | GTGTCCTTTTCTGTTTTTGG | TTTAGGCTCTCCAAACACAC | S2 or N2 |
| *PIN4* | Promoter | TACTCCTGCTTCCCTTTGTCTG | CCTTCCAGTTATCCCATCCTCT | N12 |
| *POGK* | Gene body | CCAAAAATCAGGGTATAAGCCA | GAAAAAAGCAAGGAAGATGGAA | S1 or N3 |
| *PRPF38B* | Terminal | TCGTGATAGGAAAGGGGATAGA | TACAAAAACTGGATGGAGGATG | N7 |
| *RPL13A* | Gene body | GTCCTTTTGCCCTTGTCTCC | TAGCACAGCCCATTCTACCC | S1 or N2 |
| *RPL4* | Promoter | GTGAAAGAAAAAGTGGGGCATT | ACTATTTTGACCACCCCCTACA | N7 |
| *RPS24* | Terminal | TTGTTCTCTTCATCCAGCCTTT | GGCTACTGGCTGAAAGACTACAA | N12 |
| *SNRPN70* | Terminal | TGTTTCCCCATCAGAACCAT | TTGCCTATTGTCCAGCCTTT | S2 or N1 |
| *TADA1* | Terminal | GCAGTTTCCTCTGTGCTGTATG | CCTGTTGCTTCTGTGCTATGTC | S1 or N4 |
| *TOMM20* | Terminal | TCAACTCAGCCTTTCAACACAC | AATGTCAGAATGGGTGAACTTT | N9 |
| *TOMM40* | Terminal | TGTCCATCCCCTCTTATTTTTC | ATACAGACACCCTCCTCCATTC | N8 |
| *TRIM33* | Terminal | TGTTCCTTCTCAATCTTTGG | TCAAAAATTGAGCAGAGTCC | S1 or N2 |
| *WDR3* | Promoter | GCTCAAGAATAAAAGTAGCAGAAATG | CTTGGGTGCCTTGGTTCTACT | N9 |

| Restriction Enzyme Key | | | |
|---|---|---|---|
| S1 | BamHI | N8 | HindIII-HF, SpeI |
| S2 | SpeI+NcoI | N9 | NcoI, PstI-HF |
| N1 | BamHI+NcoI+HindIII | N11 | MfeI |
| N2 | PsiI+BamHI | N12 | PsiI |
| N3 | SspI+PstI | N13 | BmtI, PvuII |
| N4 | NdeI+SpeI+NcoI | N14 | BmtI, KpnI |
| N5 | PsiI+ScaI+SspI | N15 | BsrGI, ScaI |
| N6 | SpeI+PvuII+KpnI | N17 | BsrGI, KpnI |

**Table S2:** List of footprinted loci with primers and restriction enzymes used for genome fragmentation.
